# Expecting social punishment facilitates control over a decision under uncertainty by recruiting medial prefrontal cortex

**DOI:** 10.1101/838037

**Authors:** Jaejoong Kim, Bumseok Jeong

**Author notes:** **Correspondence:** Bumseok Jeong, M.D., Ph.D., Associate Professor, Director, Laboratory of Computational Affective Neuroscience and Development, Graduate School of Medical Science and Engineering, Korea Advanced Institute of Science and Technology, Daehak-ro 291, Daejeon, Korea. ORCID ID: Jaejoong Kim, Bumseok Jeong.

## Abstract

In many decision-making situations, uncertainty facilitates suboptimal choices. However, when individuals are in a socially dangerous situation such that wrong choice would lead to a social punishment such as blame of the supervisor, they might try to minimize suboptimal choices to avoid it. In this functional MRI study, 46 participants performed a choice task in which the probability of a correct choice with a given cue and the conditional probability of blame feedback (by making an incorrect choice) changed continuously. Using computational models of behavior, we found that participants optimized their decision by suppressing the decision noise induced by uncertainty. Simultaneously, expecting blame significantly deteriorated participants’ mood. Model-based fMRI analyses and dynamic causal modeling revealed that the optimization mechanism based on the expectation of being blamed was controlled by a neural circuit centered on right medial prefrontal cortex. These results show novel behavioral and neural mechanisms regarding how humans optimize uncertain decisions under the expectation of being blamed that negatively influences mood.

**Significance Statement:** People occasionally encounter a situation that forces us to make an optimal decision under uncertainty, which is difficult, and a failure to make a good choice might be blamed by their supervisor. Although it might be hard to make right decision, they make more effort to make a good decision, which might help them to escape from the aversive outcome. However, such kind of stressful situation influences our mood to be negative. Using the computational modelling, we showed that participants computed how it is likely to be blamed and this computation motivated people to control uncertainty-induced decision noise by recruiting a neural circuit centered on the medial prefrontal cortex. However, an expectation of being blamed significantly deteriorated participants’ mood.

## Introduction

In our workplace, we make many decisions between options with uncertain values. If we fail to make a good decision, we might face a socially undesirable situation - such as being blamed by a boss. Therefore, although it could be hard to make a good decision because of an uncertainty, we may become more deliberate and expend more effort to enhance the probability of optimal choice in this situation. We occasionally encounter this kind of stressful situation which might enhance decision performance at that moment but would make us to feel bad.

The motivation to avoid negative outcomes might enhance task performance through several mechanisms, including increasing attention to the task (Engelmann, Damaraju, Padmala, & Pessoa, 2009) and enhancing working memory function (Krawczyk & D’esposito, 2013). However, the behavioral and neural mechanisms that an agent uses to optimize an uncertain decision-making process to avoid a highly probable social punishment such as blame if their decision is wrong and how such kind of social stressors influences our mood have not been investigated.

One possible mechanism of behavior under uncertainty that could be controlled under threat to optimize behavior is an exploration, which is the choice of an option that does not have maximum value among all options in the current state (Daw, O’doherty, Dayan, Seymour, & Dolan, 2006; Payzan-LeNestour & Bossaerts, 2011). More specifically, an exploration is either ‘directed’ to more uncertain options or ‘random’, which depends on decision noise induced by total uncertainty (Wilson, Geana, White, Ludvig, & Cohen, 2014) (Gershman, 2018). Because both exploration strategies help an individual obtain information about the environment, it might also benefit a long-term cumulative reward in some situations (Agrawal & Goyal, 2012). Usually, an optimal solution for the explore-exploit dilemma is difficult to compute (Gershman, 2018). However, if the uncertainties of every option are the same and an agent knows the outcomes of both selected and unselected options, an exploration would be unnecessary because it does not maximize information gain or the reward; thus, choosing the option with the maximum value would be an optimal choice. We confined the situation regarding uncertain decisions to this type of situation to simplify our question about optimizing an uncertain decision under threat. In this situation, because uncertainties among options are the same, an uncertainty-directed exploration would not exist. However, an increase in total uncertainty would increase the decision noise, causing random exploration, which is a suboptimal choice in this case that might decrease the accuracy of the choice. Therefore, we hypothesized that controlling the suboptimal choice (random exploration) driven by uncertainty-induced decision noise (UDN) might help an agent make a more accurate choice when the wrong choice is likely to result in undesirable blame. Furthermore, we expected that blame in this situation would influence not only behavior control under uncertainty but also people’s negative mood.

To test this hypothesis, we designed a task with a choice between two options whose uncertainty changes during the task and both the outcomes of selected and unselected options are fully knowable (Fig 1A). Importantly, participants received blame with high or low probabilities when they made a wrong choice, and this conditional probability changed between blocks of trials. Note that, we used blame as an aversive stimulus that motivates one to avoid it which is similar to a physical pain but encounters more frequently in our social life. However, our aim is not to compare both kind of painful stimuli in this study. We predicted that participants would infer how likely they would be blamed by making a wrong choice and would control UDN to make a more accurate choice when blame was highly likely while their mood is negatively influenced. Note that the term ‘uncertainty’ used in our study indicates an ‘estimation uncertainty’ or ‘information uncertainty’ resulting from an insufficient estimation of the value (Payzan-LeNestour & Bossaerts, 2011). In our case, uncertainty is derived from an imperfect estimation of the probabilistic association between a cue and “correct” outcome (De Berker et al., 2016). Theses hypothesis were tested by modeling participants’ behavior and mood during the task using the novel hierarchal Bayesian reinforcement learning model where an agent infers both about 1) which option is likely to be correct and how such belief is uncertain and 2) how it is likely to be blamed if one makes wrong decision and these two kind of inferences jointly influences the decision.

**Fig 1.**
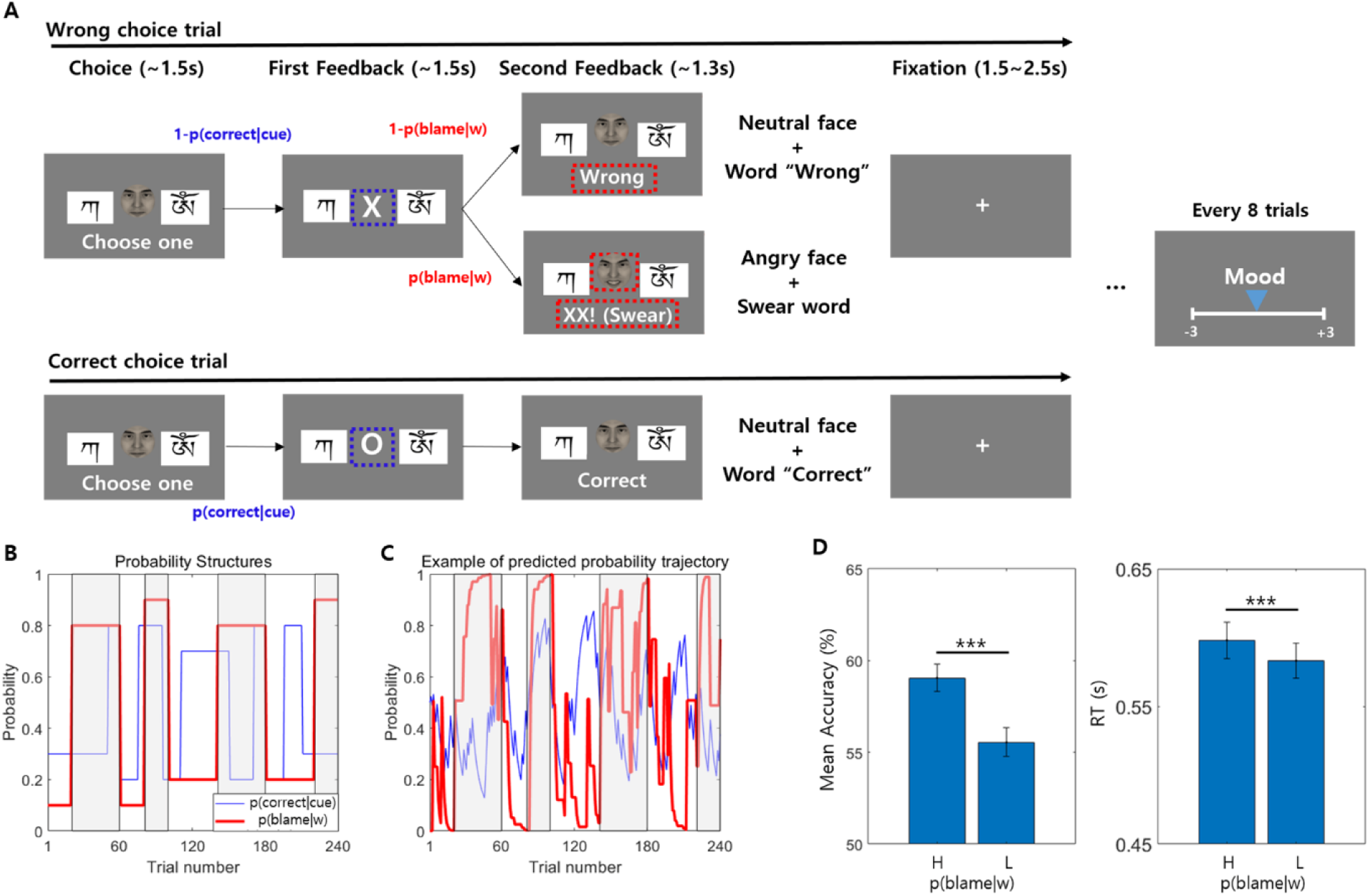
**(A) Experimental paradigm.** Two Tibetan character cues were presented, and cues were associated with a “correct” outcome with the probability of p and 1-p each. This probability p changes between blocks of trials. If participants made a wrong choice, a blame composed of an angry face and swear words appeared with the probability p(blame|w), and neutral “wrong” feedback appeared with probability 1 – p(blame|w). **(B) Probability structure of the task.** p (correct|cue) was changed between blocks of trials to change the participants’ estimation of uncertainty about this factor. Furthermore, there were high-blame blocks (gray-shaded area), and low-blame blocks. **(C) Example trajectory of p (correct|cue) and p (blame|w) estimated from the CUDN model.** This trajectory plot shows the trajectory of p(correct|cue) (blue) and p(blame|w) (red) estimated from the CUDN model in one participant, which is the computational model of behavior selected from Bayesian model selection. The plot shows the similarity between the estimated trajectory and the designed probability of both p(correct|cue) (blue) and p(blame|w). **(D) Mean accuracy difference and RT difference between blocks with high p(blame|w) and low p(blame|w).** We compared the mean accuracy and RT between high-blame and low-blame blocks using a paired t-test. Participants made more accurate choices in the high-blame blocks (t[45] = 4.09, p < 0.001), and the RT was longer within the high-blame blocks (t[45] = 4.66, p < 0.001), suggesting more deliberate and accurate choices within the high-blame blocks. The face used in this figure is different from the one that was used in the experiment to avoid using real human image in the figure (the face in this figure was generated by FaceGen Modeller (http://www.facegen.com/)).

In order to identify a neural mechanism of such behavioral optimization under threat, a model-based fMRI analyses and dynamic causal modeling (DCM) analysis (K. J. Friston, Harrison, & Penny, 2003) were performed. A recent study suggested the medial prefrontal cortex (mPFC) as a candidate region for controlling strategic avoidance behavior under threat (Mobbs et al., 2007; Qi et al., 2018). Furthermore, the mPFC implements slower, more controlled, and deliberate decision making during difficult choices (Cavanagh et al., 2011) and drives strategy shifts (Schuck et al., 2015). Therefore, we hypothesized that the neural circuit involving the mPFC would control the behavioral optimization process under the expectation of being blamed. Especially, we expected that such neural circuit would also involve rostrolateral prefrontal cortex (rlPFC) which is related to an uncertainty-driven exploration.

## Results

### Deliberate choice behavior induced by an expectation of being blamed facilitates an accurate choice

Every participant (N = 46) completed 240 trials of a choice task in which the goal was to acquire the “correct” outcome as many times as possible. In every trial, two Tibetan character cues were presented, and participants were asked to choose one (Fig 1A). Cues were probabilistically associated with either a “correct” or “wrong” outcome, and this probability was reciprocal such that

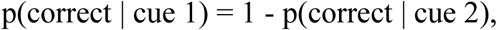

which was explicitly conveyed to participants to make the uncertainties about the probabilities of the correct choice for both cues to be equal, thus minimizing directed exploration. This probability was changed continuously to change the participants’ estimation of the uncertainty of association probability between the cue and correct choice (Fig 1B). Importantly, if the participant made a wrong choice, they were probabilistically being blamed with an angry face and swear words. We called this type of feedback blame, and we instructed participants to regard this feedback as blame in the context of social situation, such as a blame given by their boss or superior (e.g., “Please try to imagine as vividly as possible that your boss yells at you because you made a wrong choice”). Participants sufficiently practiced imagining this situation before starting the experiment (Fig 1A). Importantly, we varied the conditional probability of the appearance of the blame feedback when participants made an incorrect choice block by block with a range from 20 to 40 trials. Thus, during some blocks of the task, the conditional probability of blame feedback was very high when the choice was wrong (80% or 90%; we designated these blocks “high-blame” blocks, gray-shaded area of Fig 1B and C), while in the other blocks, this conditional probability was low (10% or 20%; we designated these blocks “low-blame” blocks). Additionally, to assess a mood of each trial, participants were instructed to rate their mood on a Likert scale ranging from −3 to 3 every eight trials after receiving secondary feedback. A total of 30 ratings were acquired from each participant. We expected that participants would implicitly or explicitly calculate the conditional probability of being blamed by making a wrong choice (we represent this subjective conditional probability of being blamed for a wrong choice as “p(blame|w)”), and this calculation would influence participants’ decision making, such as encouraging participants to make more deliberate and careful choices and mood, which was revealed in the results of the postexperimental survey (S1 Table, S2 Text).

Furthermore, the result of behavioral analyses without using computational model showed a significantly increased choice accuracy in the high-blame blocks (mean accuracy = 0.591 vs. 0.555, t[45] = 4.09, p < 0.001, CIs: 0.02 to 0.05, d = 0.6; two-tailed paired t-test; Fig 1D, left panel). Moreover, the mean response times (RTs) of the high-blame blocks were significantly longer than those of the low-blame blocks (mean RT = 598 ms vs. 583 ms, t[45] = 4.66, p < 0.001, CIs: 8.45 to 21.31, d = 0.69; two-tailed paired t-test; Fig 1D, right panel). Based on these results, participants made more accurate decisions via deliberation when p(blame|w) was high.

Note that CIs represents 95% confidence intervals, and d represents Cohen’s d. Additionally, we showed that the frequently changing probability design of p(correct|cue), which was intended to change the uncertainty of participants, influenced the choice of an option that was designed to have a lower probability of being the correct choice than the other option (i.e., a “bad” choice in our task design; beta = 0.73, t[10902] = 25.98, p < 0.001; mixed-effect logistic regression, S3 Text). This influence of probability change on a bad choice decreased in the high-blame trials (beta = −0.13, t[10902] = 4.62, p < 0.001; S3 Text), showing that participants’ choices of the bad option was influenced by our task design, which induced uncertainty and blame expectation. Note that a bad choice, which was defined with respect to the task design, differs from a suboptimal choice, which was defined by the participants’ beliefs about p(correct|cue).

### Uncertainty drives suboptimal choices, and p(blame|w) decreases uncertainty-driven suboptimal choices

In the next step, we tested whether and how uncertainty and p(blame|w) influence a suboptimal choice behavior more directly. Note that a suboptimal choice of the participant was defined as choosing an option that has a lower probability of being correct (p(correct|cue)) than other options based on their internal models or beliefs, which differed from a bad choice. In particular, we hypothesized that p(blame|w) would influence suboptimal choice behavior by controlling the degree of UDN. However, we also considered the possibility that p(blame|w) influenced suboptimal choices independent of uncertainty. Therefore, we examined 1) whether uncertainty and p(blame|w) influenced suboptimal choice and 2) whether p(blame|w) influenced this uncertainty-driven suboptimal choice. We performed a mixed-effects logistic regression analysis to explain suboptimal choice using an uncertainty, p(blame|w), and the difference in the value between two options (DV, difference in the p(correct|cue) between two cues), which also influences suboptimal choices (De Martino, Fleming, Garrett, & Dolan, 2013) and their interaction. The results showed that uncertainty significantly increased suboptimal choice behavior (beta = 0.07, t[10902] = 2.17, p = 0.03, Table 1), which likely reflects the effect of UDN. Moreover, an interaction between p(blame|w) and uncertainty significantly decreased the number of suboptimal choices (beta = −0.09, t [10902] = −3.58, p < 0.001, Table 1), whereas p(blame|w) did not influence suboptimal choice behavior (beta = −0.02, t[10902] = −0.64, p = 0.521, Table 1). Thus, p(blame|w) suppresses uncertainty-driven suboptimal choices but does not influence suboptimal choices independently of uncertainty, which is consistent with our hypothesis. Additionally, DV significantly decreased the number of suboptimal choices (beta = −0.83, t[10902] = −28.60, p < 0.001, Table 1), and the interaction between DV and uncertainty significantly increased the number of suboptimal choices (beta = 0.09, t[10902] = 3.52, p < 0.001, Table 1), suggesting that uncertainty might decrease the effect of DV on suboptimal choice. Finally, the interaction between DV and p(blame|w) was not significant (beta = −0.04, t[10902] = −1.39, p = 0.164, Table 1).

**Table 1.** The results of mixed-effects logistic regression of suboptimal choice.

| **Formula:** Suboptimal choice ~ Uncertainty + $p(\text{blame}\vert \text{w})$ + DV +<br>Uncertainty*$p(\text{blame}\vert \text{w})$ + Uncertainty*D<sub>V</sub> + D<sub>V</sub>*$p(\text{blame}\vert \text{w})$ + (1\|subject) + (1\|sex) | |
| --- | --- |
| **Fixed effect variables** | **Coefficient** |
| **Uncertainty** | $\beta = 0.07$ ($t[10902] = 2.17$, $p = 0.030$) |
| **$p(\text{blame}\vert \text{w})$** | $\beta = -0.02$ ($t[10902] = -0.64$, $p = 0.521$) |
| **DV** | $\beta = -0.83$ ($t[10902] = -28.60$, $p < 0.001$) |
| **Uncertainty\*$p(\text{blame}\vert \text{w})$** | $\beta = -0.09$ ($t[10902] = -3.58$, $p < 0.001$) |
| **Uncertainty\*D<sub>V</sub>** | $\beta = 0.09$ ($t[10902] = 3.52$, $p < 0.001$) |
| **D<sub>V</sub>\*$p(\text{blame}\vert \text{w})$** | $\beta = -0.04$ ($t[10902] = -1.39$, $p = 0.164$) |

### Computational modeling reveals that the flexible suppression of uncertainty-induced decision noise enables accurate decision making under the expectation of being blamed

Based on the results of the mixed-effects logistic regression analysis of suboptimal choices, we inferred that uncertainty increased the suboptimal choices, that p(blame|w) decreased the uncertainty-induced suboptimal choices, and that p(blame|w) did not independently influence suboptimal choices. We designed a computational model to explain these behavioral patterns such that uncertainty increases the decision noise and the p(blame|w) decreases this uncertainty-induced decision noise. This model was named as a control of uncertainty-induced decision noise (CUDN) model (*Materials and Methods*). In this model, the parameter *v* determined the degree of suppression of UDN as p(blame|w) increases. Therefore, if *v* is large for some participants, those participants greatly suppress UDN when p(blame|w) is high compared to when p(blame|w) is low and vice versa. We fitted the CUDN model to the participants’ responses and compared this model with seven other models (Table 2), including three models that do not consider the influence of p(blame|w) (RWS, HGF3S and UDN models, S4 Text), three models in which p(blame|w) has a negative value (RWS-N, HGF3S-N, UDN-N models, S4 Text) and a model in which p(blame|w) both controls the UDN and contribute as a negative value (CUDN-N, S4 Text), with random-effect Bayesian model selection (RFX-BMS) (Daunizeau, Adam, & Rigoux, 2014; Stephan, Penny, Daunizeau, Moran, & Friston, 2009). A brief summary of the eight models is provided in Table 2. Notably, in the models in which p(blame|w) has a negative value, p(blame|w) also decreased the decision noise, regardless of the current uncertainty level (uncertainty-independent control of decision noise), which was different from the CUDN model (S4 Text). The CUDN model was selected with a protected exceedance probability (PEP) of 1, suggesting that the CUDN model considering the control of UDN by p(blame|w) explains participants’ behaviors better than other models (Fig 2C) and the model attribution calculation for each participant showed that the CUDN model was the best model for 36 of the 46 participants, whereas the UDN-N model was the best for the other 10 participants (S4 Table). Furthermore, we expected that if the suppression of UDN by p(blame|w) facilitates more accurate choices in high-blame blocks, participants with a large value for the parameter *v* would make more accurate choices than participants with small or negative values for *v*. Indeed, the increase in the accuracy of choices made by participants in high-blame blocks was significantly correlated with *v* (Spearman’s ρ = 0.306, p = 0.016, Fig 2E). Furthermore, only individuals with a positive *v* showed a significant increase in accuracy (t[22] = 4.84, p < 0.001, CIs: 0.03 to 0.08, *d* = 1, using a two-tailed paired *t*-test of choice accuracy between high- and low-blame blocks), while individuals with a negative *v* did not show increased accuracy (t[22] = 1.34, p = 0.195, CIs: − 0.01 to 0.04, *d* = 0.3, using a two-tailed paired *t*-test). This result indicates that only participants who suppressed UDN exhibited increased accuracy in high-blame blocks, while others who failed to suppress UDN did not display increased accuracy.

**Table 2.** Characteristics of the computational models of behavior.

|  | **Perceptual model** |  | **Response model** |  |  |
| --- | --- | --- | --- | --- | --- |
| **Related function** | **Uncertainty calculation** | **p(blame\|w) calculation** | **p(blame\|w) contribute as a negative value** | **Uncertainty increases decision noise (UDN)** | **p(blame\|w) controls UDN** |
| **RWS** | x | x | x | x | x |
| **HGF3S** | o | x | x | x | x |
| **UDN** | o | x | x | o | x |
| **RWS-N** | x | o | o | x | x |
| **HGF3S-N** | o | x | o | x | x |
| **UDN-N** | o | o | o | o | x |
| **CUDN-N** | o | o | o | o | o |
| **CUDN** | o | o | x | o | o |

**Fig 2.**
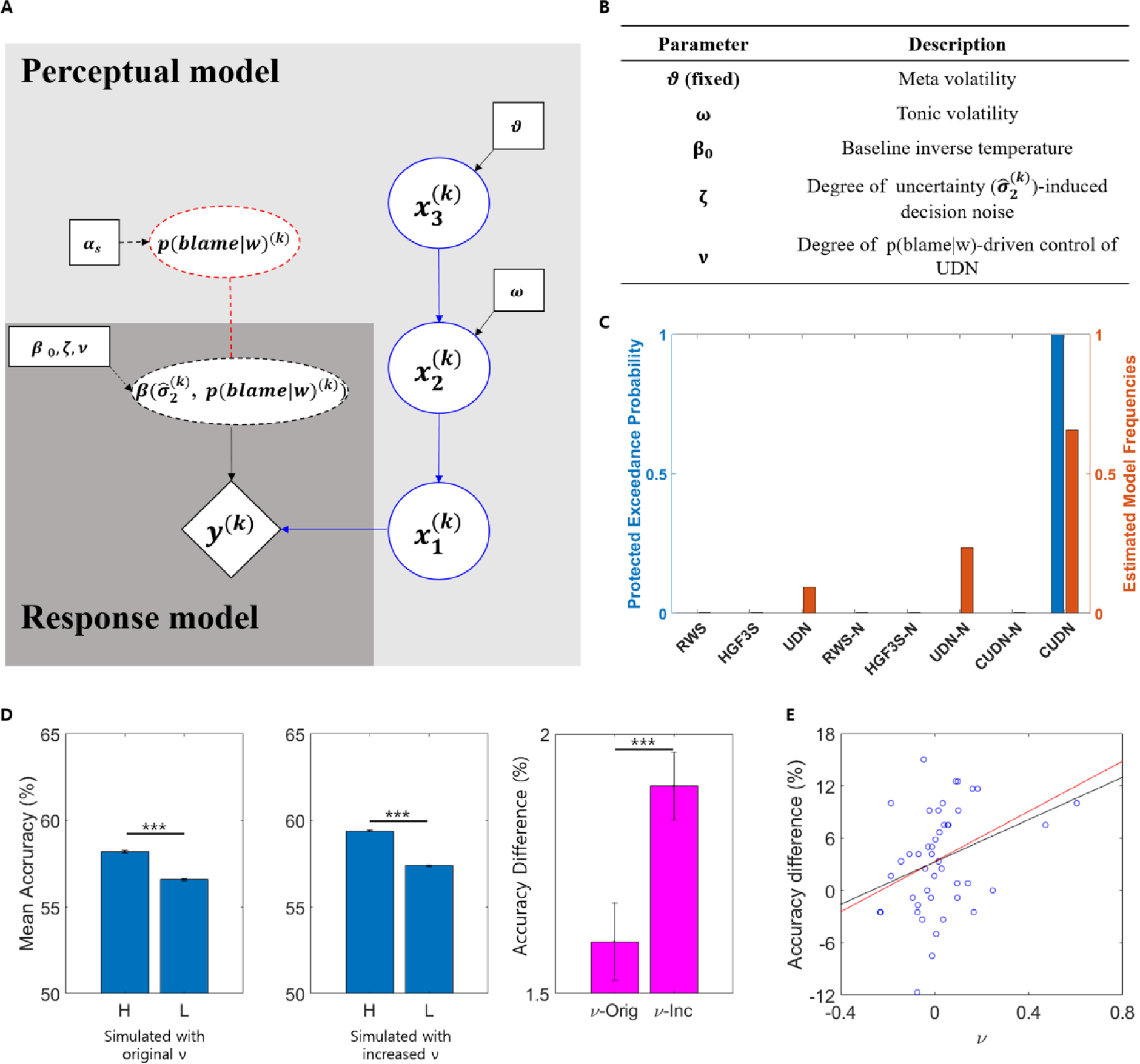
**(A) Graphic description of the computational model of behavioral optimization when expecting blame.** The CUDN model was composed of the following parts: the perceptual model and response model. The perceptual model is composed of two parallel learning systems—learning p(correct|cue) by the three-level hierarchical Gaussian filter (HGF) model (blue) and learning p(blame|w) by the Rescorla-Wagner (RW) model (red dashed circle). Importantly, in the response model, *y*^(*k*)^ represents a choice at trial k, and the inverse temperature of the Softmax function 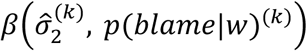 is a function of an estimation uncertainty 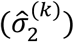 and *p*(*blame*|*w*)^(*k*)^ such that 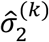 induces decision noise and *p*(*blame*|*w*)^(*k*)^ suppresses UDN. In this graphic representation, a deterministic node and relationship are represented as dashed circles and dashed lines, respectively, while solid circles and lines represent a stochastic node and relationship. **(B) Parameters of the CUDN model.** In the function 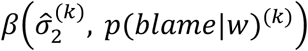, parameter *ζ* determines the participants’ degree of UDN, and *v* determines the participants’ degree of suppression on UDN as p(blame|w) increases. **(C) Bayesian model selection results.** The results of the random-effects Bayesian model selection show that the CUDN model fits the participants’ behavior better than other models (protected exceedance probability = 1). **(D) CUDN model simulation results.** Using the estimated parameters of the CUDN model for each participant, we ran a simulation 100 times per participant. We found that simulated responses recovered significantly increased accuracy in high-blame blocks (t[4599] = 21.23, left, p < 0.001). Moreover, when we increased the parameter *v* that controls the size and direction of the control of UDN, accuracy enhancement in the high-blame blocks was significantly higher than that in the simulation using the original parameter *v* (t[4599] = 3.85, right, p < 0.001). **(E) Correlation between accuracy enhancement and *v*.** The results of correlation analysis showed that the participants with large *v* showed more accuracy enhancement in high-blame blocks (Spearman’s ρ = 0.306, p = 0.016, red line). Note that this significant correlation also existed after removing points with values outside of 2 SD (rightmost 2 points) from the mean (p < 0.05, black line).

### Expectation of being blamed negatively influences mood

In order to find out whether an expectation of being blamed that influenced participants’ decision under uncertainty also influenced their mood, we performed a mixed-effect logistic regression analysis to explain participants’ moods during the experiment using p(blame|w), DV, uncertainty regarding the upcoming trial calculated from the CUDN model and the interactions between pairs of variables as fixed effects and participant and sex as random effects. The model variables used here are the same as those used in the mixed-effect logistic regression of explorative choice. Notably, p(blame|w) significantly decreased the mood of participants (beta = −0.08, t[1373] = −2.99, p = 0.003), while DV and the interaction between uncertainty and DV increased the mood of participants (beta = 0.15, t[1373] = 5.10, p < 0.001, beta = 0.09; t[1373] = 3.72, p < 0.001, respectively). No other factor influenced mood, including uncertainty (all p > 0.05).

### Model-based fMRI analysis

#### The bilateral mPFC is recruited by an expectation of blame, and the bilateral rlPFC is involved in processing a suboptimal value

In the parametric modulation analyses, robust activation of the bilateral medial prefrontal cortex (mPFC) cluster centered at [−8, 62, 28] and [6, 60, 32] (cluster-level FWE-corrected p < 0.001), the PCC cluster ([−4, −46, 34], cluster-level FWE-corrected p < 0.001) and the bilateral temporal pole cluster extending to the bilateral amygdala and hippocampus ([33, 22, −26] in the right temporal pole cluster and [−44, 16, −18] in the right temporal pole cluster, cluster-level FWE-corrected p < 0.001 in both clusters) by p(blame|w) was observed (Fig 3A and Table 3). Finally, the regions activated by the suboptimal value included the bilateral rostrolateral prefrontal cortex (rlPFC; clusters [36, 62, −8] in the right rlPFC cluster and [−18, 58, −14] in the left rlPFC cluster, cluster-level FWE-corrected p < 0.001 in both clusters, Fig 4B, S2 Fig, and S6 Table), which were involved in uncertainty-driven exploration in a previous study (Badre, Doll, Long, & Frank, 2012). Our report of the parametric modulation analysis was based on a cluster-defining threshold (CDT) of p < 0.001 and a cluster-forming size of 100 voxels. A more detailed description of the regions activated by p(blame|w), blame prediction error (BPE) (Fig 3B, *Materials and Methods*) and the suboptimal value is provided in the S5 Text.

**Table 3.** Regions activated by p(blame|w)

| **Cluster** | **Cluster p-value (FWE corrected)** | **No. of voxels** | **Region name (AAL)** | **MNI coordinates (x, y, z)** | **Voxel Z-Value (uncorrected p-value)** |
| --- | --- | --- | --- | --- | --- |
| **mPFC** | < 0.001 | 4556 | Frontal_Sup_Medial_L | -8, 62, 28 | 6.65 (<0.001) |
|  |  |  | Frontal_Sup_Medial_R | 6, 60, 32 | 5.27 (<0.001) |
| **Right TP/hippocampus/amygdala** | < 0.001 | 1156 | Temporal_Pole_Mid_R | -4, -46, 35 | 5.51 (<0.001) |
|  |  |  | Hippocampus_R | 20, 4, 18 | 4.93 (<0.001) |
| **Left TP/hippocampus/amygdala** | < 0.001 | 1368 | Temporal_Pole_Sup_L | -44, 16, -40 | 4.95 (<0.001) |
| **PCC** | < 0.001 | 407 | Cingulate_Mid_L | -4, -46, 34 | 4.61 (<0.001) |
|  |  |  | Cingulate_Post_L | -12, -46, 30 | 4.50 (<0.001) |
| **Right MTG/STG** | < 0.001 | 1120 | Temporal_Mid_R | 52, -18, -10 | 5.47 (<0.001) |
| **Left MTG/STG** | < 0.001 | 814 | Temporal_Mid_L | -56, -28, -6 | 5.13 (<0.001) |
| **Left MTG** | 0.003 | 210 | Temporal_Mid_L | -42, -60, 20 | 4.05 (<0.001) |
| **MCC** | 0.007 | 178 | Cingulate_Mid_L | -2, -18, 34 | 4.35 (<0.001) |
| **Cerebellum** | <0.001 | 817 | Cerebellum_Crus2_R | 24, -86, -36 | 5.31 (<0.001) |
| **Cerebellum** | 0.009 | 166 | Cerebellum_Crus1_L | -42, -68, -24 | 4.76 (<0.001) |
| **Cerebellum** | 0.012 | 157 | Cerebellum_Crus2_L | -22, -86, -34 | 4.66 (<0.001) |

**Fig 3.**
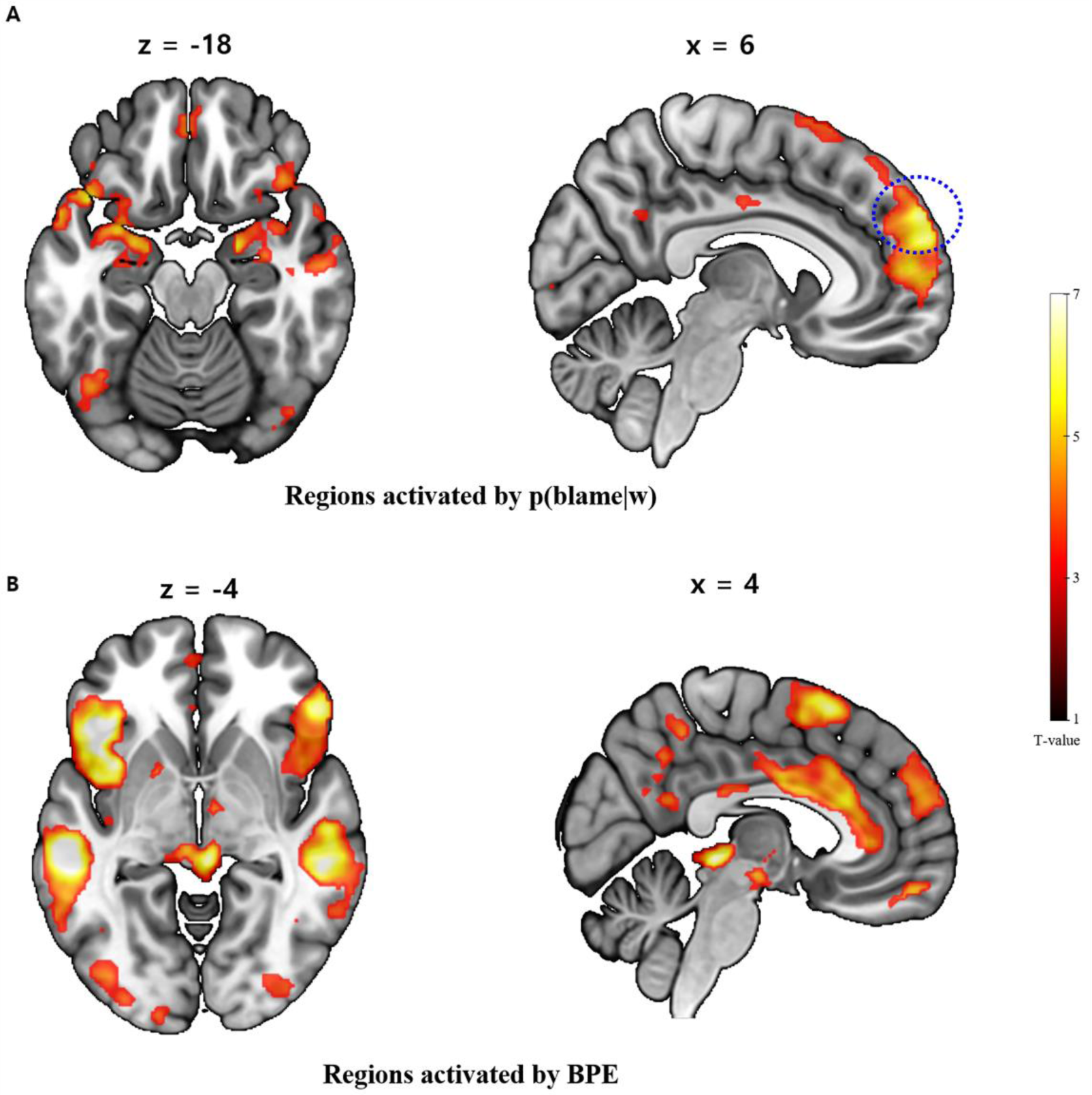
**(A) Results of the parametric modulation analysis by p(blame|w).** MPFC, PCC (center) and hippocampus (left), were activated by p(blame|w) (all cluster-level FWE-corrected p < 0.001). The blue dashed circle denotes the right mPFC used in the DCM as a region of interest (ROI). **(B) Results of the parametric modulation analysis by blame prediction error (BPE).** In the parametric modulation analysis of BPE, periaqueductal gray matter (PAG) and anterior/middle cingulate cortex (ACC/MCC) clusters appeared. Moreover, regions similar to p(blame|w), including the mPFC, appeared (all cluster-level FWE-corrected p < 0.001).

**Fig 4.**
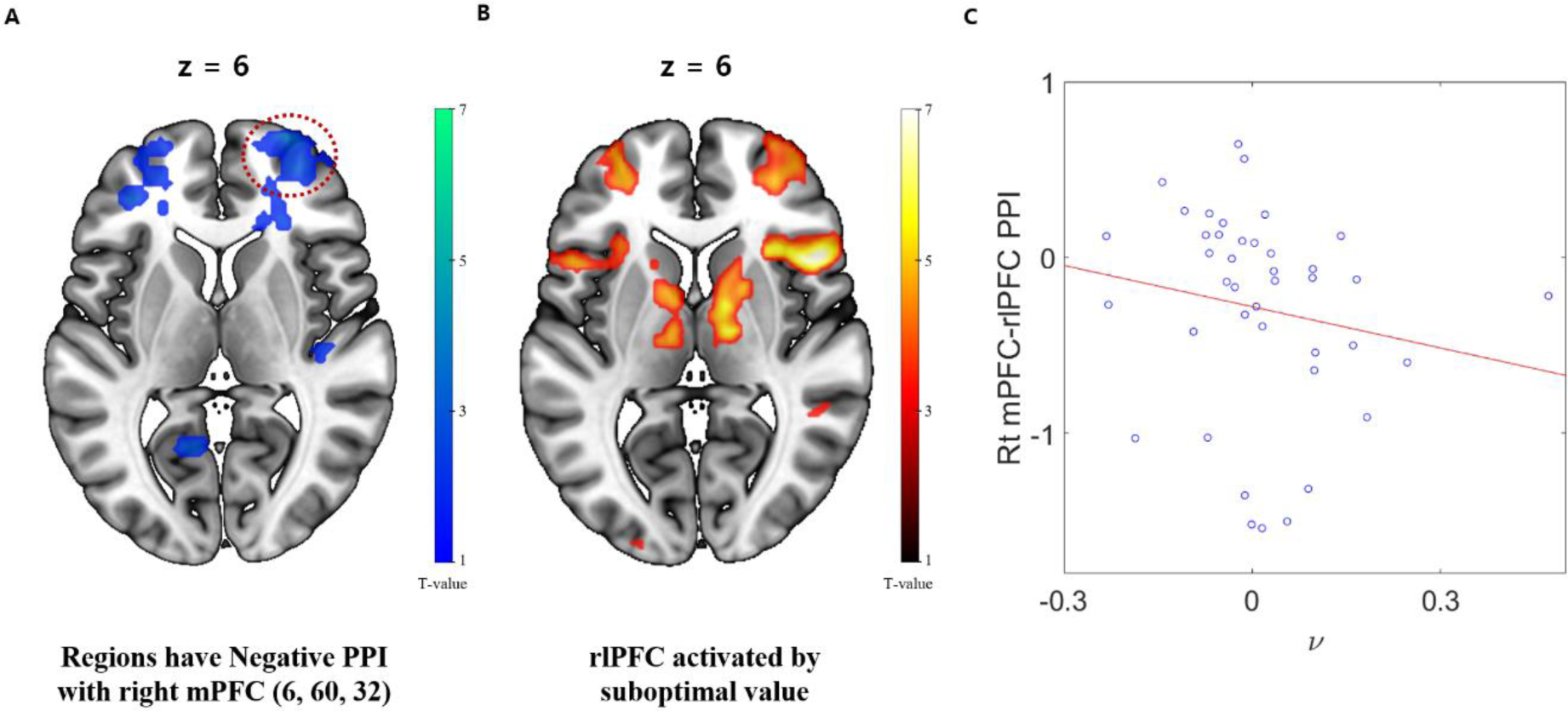
**(A) The PPI results of the right mPFC BOLD time series as a physiological factor and p(blame|w) as a psychological factor.** The results of the PPI analysis showed that functional connectivity between the right mPFC and right rlPFC was negatively modulated as p(blame|w) increased (red dashed circle, cluster-level FWE-corrected p = 0.039). Notably, the negative PPI with the left rlPFC was marginally significant (cluster-level FWE-corrected p = 0.065). **(B) rlPFC clusters activated by the suboptimal value.** Bilateral rlPFC was also involved in the processing of the suboptimal value (both cluster-level FWE-corrected p < 0.001). (C) **Results of the correlation analysis between the right mPFC-rlPFC PPI and *v*.** There was a significant correlation between the negative modulation of right mPFC-rlPFC functional connectivity and the parameter *v* that determines the size of control on UDN by p(blame|w) (Spearman’s ρ = −0.34, p = 0.029).

#### PPI analysis reveals a correlation between UDN suppression and the negative modulation of the functional connectivity between the right mPFC and the right rlPFC

In the parametric modulation analysis, we found that the bilateral mPFC was robustly activated by p(blame|w), which is consistent with our hypothesis. We suspected that if the mPFC is involved in the suppression of UDN by the blame expectation, an interaction pattern between this region and the regions activated by the suboptimal value would change depending on the level of the p(blame|w). To test this hypothesis, we performed a PPI analysis (K. Friston et al., 1997; McLaren, Ries, Xu, & Johnson, 2012). Interestingly, there was a negative cluster in the right rlPFC showing a negative modulation of the functional connectivity with right mPFC by p(blame|w) ([28, 66, 6], cluster-level FWE-corrected p = 0.039, Fig 4A and S8 Table). Notably, this cluster exhibited a large overlap with the right rlPFC clusters activated by the suboptimal value (Fig 4A, 4B). Moreover, negative modulation of the functional connectivity between the right rlPFC and right mPFC was significantly correlated with *v* (Spearman’s ρ = − 0.34, p = 0.029, Fig 4C). These results support our hypothesis that the control of the UDN is related to a modulation of interaction between mPFC and the suboptimal value-related region depending on a level of the p(blame |w). Except the rlPFC cluster with right mPFC seed, three more significant clusters were found in this PPI analyses (one another cluster with the right mPFC seed and two cluster in the left mPFC seed) those did not showed a significant correlation with *v*. These results are provided in the S8 Table (S8 Table).

#### Dynamic causal modeling reveals that the suppression of UDN is related to the negative modulation of effective connectivity from the right rlPFC to the right mPFC

After identifying that negative modulation of the functional connectivity between the right mPFC and right rlPFC is related to the suppression of UDN by p(blame|w), we performed a DCM analysis to identify a neural dynamics model, including causal relationships between both regions, and to determine which part of this neural dynamics model is specifically related to the suppression of UDN by p(blame|w). In the RFX-BMS between 16 DCM models, model 7 (Fig 5), which includes a driving input to the right mPFC by p(blame|w), a bidirectional fixed connection between two regions and modulation of both connections by p(blame|w), was selected with a PEP of 0.999. We conducted a robust linear regression analysis using the parameter *v* as a dependent variable and two effective connectivities, one from the right mPFC to the right rlPFC and another from the right rlPFC to the right mPFC in model 7, as independent variables to identify which direction of modulation by p(blame|w) was related to the suppression of UDN. Only the effective connectivity from the right rlPFC to the right mPFC negatively influenced *v* (β = −0.46,t[29] = −3.09, p = 0.004, CIs: −0.75 to −0.17, *d* = −0.5), whereas the effective connectivity in the opposite direction did not (β = −0.2,t[29] = −1.33, p = 0.193, CIs: −0.49 to 0.09, *d* = −0.2). Furthermore, because the modulation from the baseline ‘fixed’ connectivity was related to the modulation of UDN by p(blame|w), we suspected that the fixed connection from the right rlPFC to the right mPFC might be related to the UDN and the modulation of this connection is related to the control of UDN. To test this hypothesis, we performed a robust linear regression analysis using the parameter ζ, which is related to the UDN as the dependent variable and the fixed connection from the right rlPFC to the right mPFC as an independent variable, which showed a positive relationship between two variables (β = 0.24, t[30] = 2.08, p = 0.047, CIs: 0.01 to 0.46, d = 0.4, Fig 5). In summary, we showed that the UDN is related to the fixed connection from the right rlPFC to the right mPFC and the modulation of this connection via p(blame|w) is related to UDN suppression.

**Fig 5.**
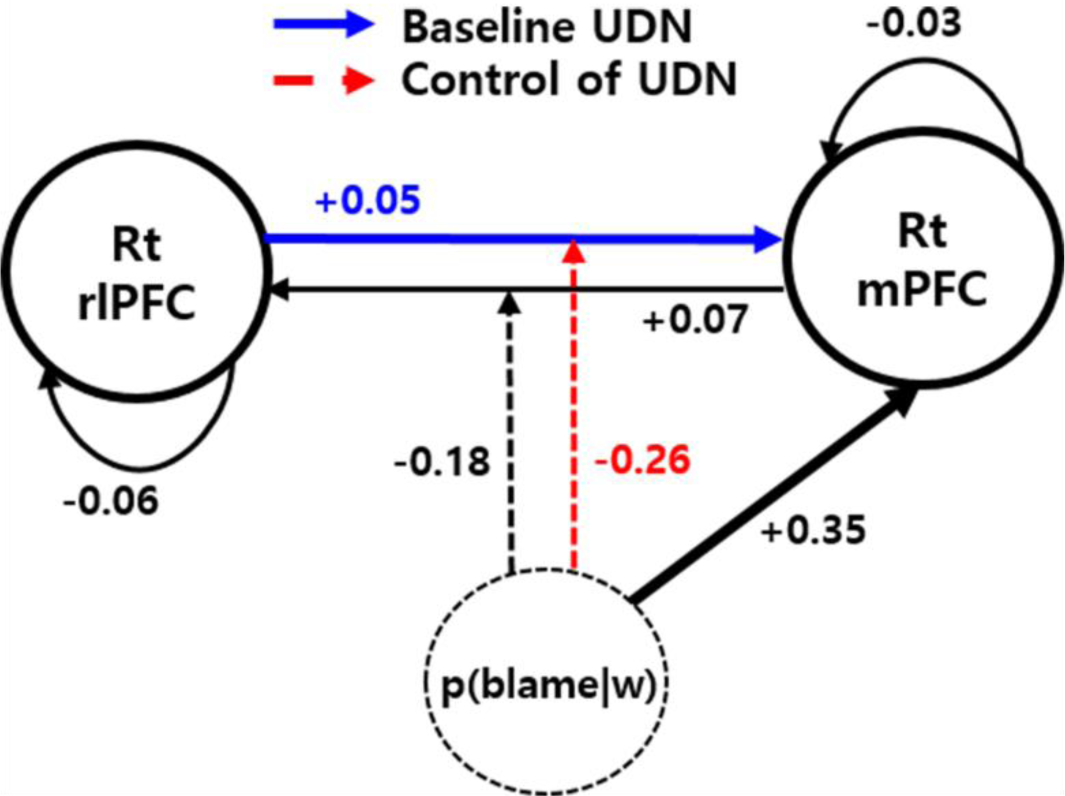
The winning model of the Bayesian model selection between DCM models. The winning model showed the bidirectional fixed connections between the right mPFC (blue dashed circle in Fig 3) and rlPFC (red dashed circle in Fig 4). Both connections were modulated by p(blame|w) and p(blame|w) also acted as a driving input to the mPFC. The fixed connection from the right rlPFC to the mPFC (blue arrow) was related to the degree of UDN. Moreover, p(blame|w) acted as a driving input to the right mPFC, and the effective connectivities from the right mPFC to the rlPFC as well as from the right rlPFC to the mPFC were modulated by p(blame|w) (red dashed line). The modulation of effective connectivity from the right rlPFC to the mPFC was to related to the degree of UDN control via p(blame|w).

## Discussion

Under the condition that a wrong decision leads to severe blame by another, we must regulate ourselves to make better decisions. In this study, participants made more accurate choices as a conditional probability of blame received by making a wrong choice through the suppression of UDN. However, expecting such blame with high probability impaired participants’ mood. Furthermore, fMRI analyses, including DCM analyses, revealed that a neural mechanism underlying this behavioral tendency is related to the suppression of connectivity from the right rlPFC to the mPFC as p(blame|w) increases, where the right rlPFC was activated by the suboptimal value and the right mPFC was activated by p(blame|w).

Our study improves our understanding of the behavioral and neural mechanisms of optimal decision-making strategies to avoid aversive outcomes expected for incorrect decisions in a social context and how such kind of stressor influences our mood. From a behavioral perspective, when punishment or loss is expected with lower performance, participants enhance their performance by enhancing working memory function (Krawczyk & D’esposito, 2013) or inhibitory control mechanisms (Simões-Franklin, Hester, Shpaner, Foxe, & Garavan, 2010). However, to the best of our knowledge, few studies have proposed the computational mechanisms that are involved in optimizing uncertain decisions under punishment expectations based on explicit computational models of behavior. In the present study, using the CUDN model, we successfully showed that participants made more accurate choices when a conditional probability of blame given a wrong choice was high through the flexible control of UDN.

In many cases, random exploration might be beneficial for maximizing a long-term expected reward because it helps the individual obtain information about uncertain options (Agrawal & Goyal, 2012; Payzan-LeNestour & Bossaerts, 2011). However, in situations such as our task, where there is no more information gain by choosing one option over another, random exploration would become a suboptimal choice that might decrease the accuracy of the choice. According to the results from our mixed-effects logistic regression analysis, even in this situation, uncertainty about the correct probability, which corresponds to total uncertainty in Gershman *et al*. (Gershman, 2018), increased the suboptimal choices, which corresponds to random exploration. Interestingly, p(blame|w) decreased these “uncertainty-driven” suboptimal choices but did not directly influence the suboptimal choice behavior.

We modeled this effect of expectation of being blamed on uncertainty-driven suboptimal choices using a CUDN model in which an uncertainty increases decision noise and p(blame|w) controls this UDN this model explained participants’ behaviors better than other models. An interesting point regarding models where p(blame|w) had a negative value is that p(blame|w) naturally decreased the decision noise, regardless of the amount of uncertainty, which is different from the CUDN model in which only UDN is influenced by p(blame|w). Therefore, another point recognized from the Bayesian model selection results is that p(blame|w) likely controls the decision noise in a manner proportional to a degree of uncertainty but not in an uncertainty-independent manner. This hypothesis is consistent with the results of the mixed-effect logistic regression analysis, where p(blame|w) decreased the uncertainty-driven suboptimal choices but did not directly influence the suboptimal choice behavior. Because the decision noise increases with uncertainty, the requirement for the suppression of UDN to make an optimal choice is higher when high uncertainty exists. Furthermore, based on the results of the post-experimental survey, we recognized that participants made more deliberate, effort-driven decisions under threat, resulting in the suppression of (uncertainty-driven) decision noise. Thus, we surmise that the suppression of decision noise might require a mental effort that is costly to exert (Botvinick & Braver, 2015; Shenhav et al., 2017). Therefore, we speculated that a balance between the cost of deliberation and the need to suppress decision noise to make an optimal choice under uncertainty might increase the efficiency of the suppression of UDN, particularly in a volatile environment in which an uncertainty level continuously changes, such as in our task.

After confirming that p(blame|w) controlled the UDN, we revealed that the suppression of UDN enabled the participant to make an accurate choice by showing a positive correlation between increased accuracy in the high-blame blocks and the model parameter *v*. Furthermore, the results of the simulations of the CUDN model confirmed this finding, as an increase in *v* increased the accuracy to a greater extent in the high-blame blocks. Therefore, our behavioral results suggest that when a wrong choice is likely to result in an aversive outcome under the uncertain decision-making situation used in our task, an agent tries to make an accurate choice by suppressing UDN to some degree. However, we also have shown that an inference regarding a conditional probability of the blame significantly impaired participants’ mood. People suffer from an abusive supervision are at high risk of affective disorder (Tepper, Moss, Lockhart, & Carr, 2007). We speculate that people who are frequently exposed to an abusive supervision similar to our task such that their supervisor forces them to make good decision under uncertainty with social punishment would be more likely to evolve an affective disorder by an accumulation of a daily negative mood induced by an aversive outcome expectation by their wrong decision.

We then identified the neural mechanism underlying the expectation of blame and the suppression of the UDN by p(blame|w). The regions involved in processing p(blame|w) included the mPFC, hippocampus and PCC, which are similar to the regions involved in the “cognitive fear circuitry” (Qi et al., 2018). The suggested role for this circuit was strategic avoidance of a threat (Qi et al., 2018) and the authors mentioned that the mPFC is likely involved in selecting defensive response strategies (Qi et al., 2018). This region contains “strategy-selective” cells that protect against threats, as reported in an animal study (Halladay & Blair, 2015). Moreover, the mPFC is related to internally driven strategy shifts (Schuck et al., 2015). Consistent with these studies, subsequent PPI and the DCM analyses revealed the involvement of the mPFC in the control of UDN. Especially, neural dynamics involving the right mPFC and rlPFC was related to the control of UDN via p(blame|w). Two regions had bilateral fixed connections, and these connections were negatively modulated according to the p(blame|w). Importantly, among these two directional connections, only the connection from the right rlPFC to the right mPFC was relevant to the control of UDN, such that the UDN itself was related to the fixed connection from the right rlPFC to the right mPFC, and the control of the UDN was related to the modulation of this connection via p(blame|w).

Based on these results, we propose a possible neural “gate” model regarding the control of the UDN based on p(blame|w) that fits with the structure of the CUDN model. In this model, during decision making, the information flow from the right rlPFC to the mPFC encodes the degree of decision noise induced by a given level of uncertainty (UDN), and this flow is regulated by the gate-like mechanism of the mPFC, where this gate is controlled by the input conveying a level of p(blame|w). Thus, the activation of the mPFC due to p(blame|w) blocks the flow from the right rlPFC to mPFC to reduce UDN. In the CUDN model, a probability of the suboptimal choice is determined by following three components. 1) After the value of options are computed, the probability of suboptimal choice increases as the suboptimal value increases. 2) Given the suboptimal value, an uncertainty induces decision noise that increases the probability of the suboptimal choice. 3) p(blame|w) reduces the UDN. Here we propose neural mechanisms those corresponds to each of three components of the CUDN model such that an activation of rlPFC by suboptimal value corresponds to the first component and the fixed connection from the right rlPFC to the mPFC, which pathway might reflect an integration of UDN to the first component, thus corresponds to the second components. Final neural mechanism that corresponds to the third component is that, the right mPFC receives the information regarding p(blame|w) which activates this region as the p(blame|w) increases and by using this information, the mPFC regulates the flow from the right rlPFC to the mPFC to control the UDN which could explain the concurrent activation of the mPFC and the negative modulation of the connection from the right rlPFC and the mPFC based on p(blame|w). This gate model not only corroborates our behavioral and fMRI analyses but also is consistent with previous studies suggesting that the mPFC controls strategic avoidance behavior under threat (Qi et al., 2018). Furthermore, a previous study suggested that the mPFC controls the slower and deliberate decision making associated with difficult choices by modifying the decision threshold of the drift-diffusion model via an interaction with the subthalamic nucleus (Cavanagh et al., 2011), which supports our gate model and suggests a possible extension of the gate model to include the subthalamic nucleus and the basal ganglial motor control system.

Finally, the PE signal produced by unexpected blame involved the PAG and bilateral ACC/MCC (S12 Text, Fig 2B, and S5 Table). In particular, the PAG signals an APE induced by physical pain (Roy et al., 2014). In this previous study, the APE signal was conveyed to the anterior MCC (aMCC) (Roy et al., 2014). Similar to the APE signal in the previous study, in the PPI analysis that used the PAG ROI, the connectivity between the PAG and aMCC area increased with BPE (S12 Text, S4 Fig, and S9 Table). Based on these results, an updating mechanism of blame expectation due to BPE is similar to the physical pain expectation mediated by the APE. This finding is consistent with previous studies showing similarities between physical and social pain (Eisenberger, 2012). However, the dmPFC, which is more related to social pain than physical pain, was also activated by BPE (Woo et al., 2014).

Our experiment used blame, which is a type of social punishment, as punishment feedback. Because an aim of our experiment was not a comparison between an effect of social and non-social punishment, we did not perform a similar experiment using a non-social punishment (e.g., physical pain or monetary loss). Therefore, it is difficult to determine whether this effect is universal to all types of punishment or specific to blame. However, the results of this study allow us to infer the similar or different effect of blame with regard to other non-social stimuli. For example, if the expectation of blame is similar to that of loss, we would expect that the p(blame|w) would contribute as a negative value; however, this result was not the case in the behavioral modeling section. Furthermore, previous studies have shown that physical and social pain are similar in both emotional response and saliency as well as share a similar neural representation (Eisenberger, 2012). Moreover, we observed a similar between updating blame expectation and updating of physical pain expectation (Roy et al., 2014). Therefore, we hypothesize that the effect of expecting blame might be more similar to that of expecting physical pain than to that of expecting monetary loss. However, this hypothesis is only speculative, and an experiment using non-social stimuli might help us to identify both the similar and different neural and behavior mechanisms that underlie the optimization of behavior under social and non-social threats.

Final notable point is that, although the DCM analysis enabled us to specify the neural dynamics related to UDN suppression, this analysis was performed only on the participants who showed the significant experimental effect of parametric modulation based on p(blame|w) and the negative PPI effect (Stephan et al., 2010). Therefore, we are not able to confirm whether the neural dynamics of the participants who did not show the experimental effect are similar to those included in the analysis.

In conclusion, we identified one strategy for optimizing uncertain decision making under a threat and the underlying neural mechanism. Because there was no benefit of the suboptimal choice in our task, the suppression of UDN when receiving blame likely increased the accuracy of participants’ decisions in those situations, and this phenomenon was successfully modeled using the CUDN model. On the other hand, an expectation of being blamed deteriorated participants’ mood. The implementation of this behavioral optimization strategy was related to the suppression of effective connectivity from the right rlPFC to the right mPFC as p(blame|w) increased. These results added one novel neural mechanism of a brain region related to processing threat that actually interacted with other decision-making-related regions to avoid a threatening outcome.

Because we addressed only one optimization mechanism under particular conditions, where directed exploration is absent or minimal, an extension of our research to determine how directed exploration is influenced in this situation would be interesting. Based on recent findings that people became more ‘myopic’ under the threat (Korn & Bach, 2019), we speculate that directed exploration would also be reduced by blame expectation and it could explain a reduced creativity under the threat. Finally, from the perspective of computational psychiatry, an investigation of the optimization behavior of our task in patients with psychiatric conditions, such as autism and psychopathologies, would be interesting to quantify their lack of an ability to expect social responses and utilize adaptive behaviors (Sevgi, Diaconescu, Tittgemeyer, & Schilbach, 2016).

## Materials and Methods

### Ethics Statement

All participants provided written informed consent to participate in the experiment based on sufficient explanation about the study (including blame). The study was approved by the KAIST Institutional Review Boards (IRB) in accordance with the Declaration of Helsinki.

### Participants

Forty-six participants (32 males and 14 females, mean age of 22.61 ± 3.61 years) from the KAIST volunteered for this experiment. A structured interview was conducted by three psychiatrists using the Mini International Neuropsychiatric Interview (MINI) web version (Sheehan et al., 1998). Additionally, participants completed psychiatric/psychologic surveys, including the Patient Health Questionnaire-9 (Kroenke, Spitzer, & Williams, 2001) and General Anxiety Disorder-7 (GAD-7) (Spitzer, Kroenke, Williams, & Löwe, 2006). Among the forty-six participants, eight participants were diagnosed with major depressive disorder (MDD, S8 Text). All forty-six participants underwent fMRI experiments on one day, and structural T1 MRI scans were acquired on another day. We failed to acquire structural T1 MRI images for three participants (YAD_10034, YAD_10039, and YAD_10055), and another participant was excluded because of asymptomatic cerebral vascular anomalies, which were detected first time during the MRI scan (YAD_10083). Finally, the fMRI data from one participant were excluded from the analysis because of excessive head movements (YAD_10008). All the experimental data, except the MRI data, from the forty-six participants were used in the behavioral analysis, and the data from forty-one participants were used in the fMRI analysis.

### Experimental task

Each participant completed 240 trials of a choice task with the goal of acquiring the “correct” outcome as many times as possible. In every trial, two Tibetan character cues were presented for 1500 ms on both sides of the screen, and participants were asked to choose one of them (Fig 1A). Each cue was probabilistically associated with either a “correct” or “wrong” outcome. This probability changed between blocks of trials with a length ranging from 10 trials to 50 trials to continuously change participants’ strength of belief about the association probability (in other words, uncertainty regarding the probability) (Fig 1B). This estimation uncertainty about the probability varies according to the volatility of the probabilistic environment (De Berker et al., 2016). For example, if the probabilistic environment changes quickly (the probability change occurs within a short period), an agent might not strongly believe their present estimate of the association probability and would be ready to change their beliefs. Participants were instructed to make decisions while considering the possibility of a cue-outcome contingency change. We also instructed participants that if the chosen option is “correct”, the other option must be “wrong”, and the sum of the probabilities of the correct choice for each cue is indicated by the following equation:

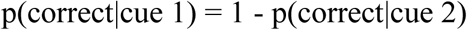

These instructions were intended to make the uncertainties about the probabilities of the correct choice for both cues equal, thus minimizing directed exploration.

Feedback consisted of two stages. The first feedback provided information about whether the choice was correct or wrong, and the second feedback provided contextual information about whether a wrong decision led to blame. More specifically, after 1500 ms of cue presentation, the first feedback appeared for 1500 ms, which was “O” for a correct choice and “X” for a wrong choice (Fig 1A). Subsequently, the secondary feedback appeared on the screen for 1300 ms. If the choice was correct, a male face with a neutral expression and the word “Correct” appeared on the screen. However, when participants made a wrong choice during that trial, one of two types of secondary feedback was presented pseudorandomly. In some trials, the secondary feedback for a wrong choice was the male face with a neutral expression and the word “Wrong”, which is similar to the secondary feedback for the correct choice, and it was called “neutral wrong feedback” (NF). However, in the other wrong trials, the secondary feedback was an angry male face with a swear word that represents a socially threatening response (e.g., “You fxxking stupid idiot!”). We called this type of feedback blame. Notably, we instructed participants to regard this feedback as given by their boss or superior (e.g., “Please try to imagine as vividly as possible that your boss swears at you because you made a wrong choice.”) to ensure that this feedback was socially threatening to every participant. Participants sufficiently practiced imagining this situation before starting the experiment (Fig 1A). Therefore, the participants’ feelings toward blame were significantly more negative than their feelings toward NF (t[44] = 16.7, p < 0.001 using a two-tailed paired *t-*test, S2 Text). In the preexperimental task exercise session, participants completed 40 trials of the task, which had a similar structure to the task of the main experiment, to familiarize them with the task. Importantly, we varied the conditional probability of the appearance of the blame feedback when participants made an incorrect choice block by block with ranges from 20 to 40 trials. Thus, during some blocks of the task, the conditional probability of blame feedback was very high when the choice was wrong (80% or 90%; we designated these blocks as “high-blame” blocks, gray-shaded area of Figs 1B and C), while in the other blocks, this conditional probability was low (10% or 20%; we designated these blocks as “low-blame” blocks). We expected that participants would implicitly or explicitly calculate the conditional probability of blame by making a wrong choice (we represent this subjective conditional probability of blame for a wrong choice as a “p(blame|w)”), and this calculation would influence participants’ decision making. The probabilistic schedule of both the cue-outcome contingency and conditional probability of blame feedback changed independently during the task. Furthermore, this probabilistic schedule was presented in the opposite order to twenty-four of the forty-six participants to avoid the effect of block order. Additionally, in every eight trials, participants were asked to rate their mood between −3 and +3 to determine the extent to which blame influenced their moods during the task. Finally, after the experiment, participants completed a post-experimental survey regarding the valence of feedback stimuli and the various effects of feedback stimuli on the tasks, including an effect on decision making and mood (S1 Table).

### Difference in choice accuracies and mean response times between blocks with high-/low-blame blocks

We first tested whether the expectation of blame for making a wrong choice increased accuracy by comparing the mean accuracies of choices between high-blame blocks and low-blame blocks using two-tailed paired *t*-tests. Both high- and low-blame blocks contained 120 trials (half of the 240 trials), and accuracy was defined as the proportion of correct responses in each block. Additionally, using two-tailed paired *t*-tests, we compared the mean response time between high- and low-blame blocks, which likely reflects the degree of deliberation.

### Mixed-effect logistic regression analysis of suboptimal choice

In the previous behavioral analysis, we showed that the likelihood of choosing the bad option was increased by the task design intended to induce an uncertainty, and this tendency decreased in trials with a high probability of blame feedback. In the next step, we tested whether and how actual uncertainty and p(blame|w) influence a suboptimal choice behavior more directly. In particular, we hypothesized that p(blame|w) would influence their suboptimal choice behavior by controlling the degree of UDN. However, we also considered another possibility: that p(blame|w) would influence suboptimal choice, regardless of uncertainty. Therefore, we examined whether 1) uncertainty and p(blame|w) influences suboptimal choice and 2) p(blame|w) influences this uncertainty-driven suboptimal choice. However, participants’ internal beliefs about p(correct|cue), its uncertainty, p(blame|w) and other decision variables cannot be observed directly; rather, they must be estimated by fitting a computational model of behavior. Furthermore, a suboptimal choice was also defined with respect to the belief of participants, i.e., the choice of an option believed to be less likely to be correct p(correct|cue) (Daw et al., 2006). To estimate these variables regarding participants’ internal beliefs, we used trial-by-trial estimates of these variables derived from computational models of the behavior determined to be optimal for estimation. Particularly, p(correct|cue) and its uncertainty was extracted by fitting the three-level hierarchical filter model with a basic Softmax response function (HGF3S) to the response of participants because this model is optimal for explaining behavior in the context of a learning task with a volatile probability structure such as ours (C. Mathys, Daunizeau, Friston, & Stephan, 2011). Furthermore, p(blame|w) was estimated using the Rescorla-Wagner (RW) model (Rescorla & Wagner, 1972), which is an optimal model for learning the Pavlovian value of a stimulus, with a fixed learning rate of 0.5 for all participants; thus, we assumed a blame expectation that changes based on the difference between current expectations and the existence of blame feedback given an incorrect choice. Descriptions of these behavioral models are provided in the next section. Using the variables extracted from the computational model, we performed a mixed-effect logistic regression analysis to explain suboptimal choice behavior. Suboptimal choice was defined as the choice of an option with a lower belief of p(correct|cue) than the other, and this choice was used as the dependent variable. An uncertainty, p(blame|w), DV, which is the difference in the value between the two options (the difference of the model-driven p(correct|cue) between two cues) (Boldt, Blundell, & De Martino, 2019), and their interactions were used as fixed effects. Subject and sex were used as random effects. Importantly, the model-fitting procedure in this analysis was performed only to extract variables for the mixed-effect logistic regression analysis and was independent of the model-fitting procedure in the following explicit computational modeling section. For example, we did not consider modeling the integrative effects of p(blame|w) or p(correct|cue) on the decision as in the CUDN model or the other models in this analysis, and the calculations of these two variables were assumed to be independent; thus, they were estimated separately. For a similar reason, a basic Softmax response model, which does not consider the effects of uncertainty or p(blame|w) on choice behavior, was used because we were unable to determine the influences of uncertainty and p(blame|w) on choice in this step (we will attempt to investigate whether they influence suboptimal choice). The mixed-effect logistic regression analysis was implemented using the MATLAB function “fitlme”.

### Behavior model of choice

Based on the results of the mixed-effects logistic regression analysis of suboptimal choice, we inferred that uncertainty increases suboptimal choices, and p(blame|w) decreases uncertainty-induced suboptimal choices but did not independently influence suboptimal choice. We designed a computational model to explain these behavioral patterns such that uncertainty increases decision noise to increase suboptimal choice, and p(blame|w) decreases this noise. This model was named the “control of uncertainty-induced decision noise” (CUDN) model. Furthermore, we tested other models that might explain this behavior. In particular, we suspected that p(blame|w) integrates as a negative decision value during decision making. Therefore, models regarding p(blame|w) as a negative value were constructed (S4 Text). Model fitting was performed using the functions of the TAPAS toolbox (http://www.translationalneuromodeling.org/tapas), and the preexisting models used in Bayesian model selection were implemented in the same toolbox with the default prior parameter settings.

The CUDN model was composed of a perceptual model and a response model (Fig 2A). In the perceptual model, the agents’ beliefs about probability and the correct choice cue were modeled using an HGF3 (C. Mathys et al., 2011), and the calculation of p(blame|w) was modeled using the RW model (Rescorla & Wagner, 1972).

In the HGF3 model, an agent is assumed to continuously update the internal generative model regarding the hidden cause of the sensory input assumed to be hierarchically structured (x_1_, x_2_, and x_3_). This model assumes that environmental hidden states evolve as a Gaussian random walk (in the case of x_2_ and x_3_) whose step size is influenced by the above level; thus, both the current belief and step size influence the predicted uncertainty of each level. Links between each level are formulated using the following equations:

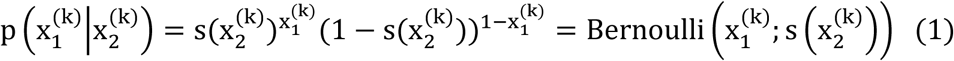

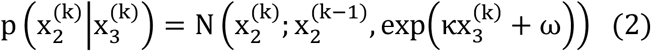

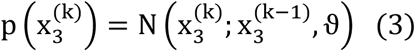

where s is the following sigmoid function:

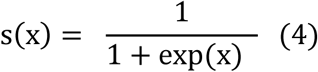

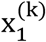 represents the prediction of a “correct” given cue that takes 0 or 1 that follows a Bernoulli distribution; 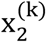 represents the inferred probability of 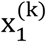 in a logit space; and 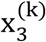 represents the inferred log volatility of the environment that reflects the changes in the correct probability of the given cue. The parameter κ describes the degree of influence on the second level based on the volatility of the third level, and we fixed this parameter to 1. The parameter ω reflects the tonic component of environmental volatility, which was estimated as a free parameter, and ϑ reflects the variance in the volatility at level 3, which is sometimes called “meta volatility”. This parameter was fixed to the default value provided by the TAPAS toolbox.

By inverting this model given sensory input, an agent updates his or her generative model as an ideal Bayesian observer. This inversion was performed using the variational Bayesian scheme, resulting in a set of closed-form update equations that provide trial-by-trial updates of the state variables, including hidden state variables such that the beliefs regarding each trial were updated by precision-weighted PE at levels 2 and 3. This idea was formulated as follows:

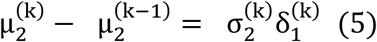

where 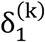 is the PE at trial k, and 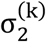 is the posterior variance at level 2

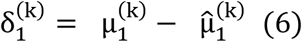

and 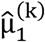 is the prediction of the correct choice resulting from a sigmoidal transformation of the previous belief about the probability of a correct choice 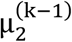

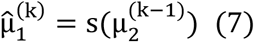

Finally, the update is proportional to the precision-weighted PE at level 3:

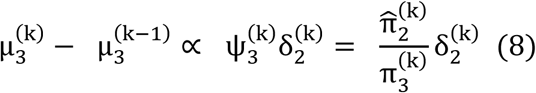

where 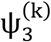 is the precision ratio, and 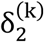 is the volatility PE formulated as

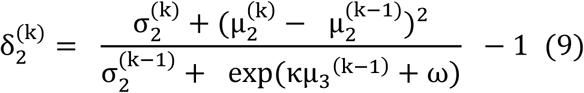

Our particular interest concerns the estimation uncertainty at the second level, representing uncertainty in the belief regarding the probability of a correct choice. In the HGF3 model, this uncertainty was calculated as follows:

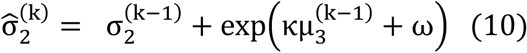

which means that the estimation uncertainty at the presentation of the cue is influenced by posterior uncertainty at the second level of the previous trial and the volatility of the third level. Therefore, as the volatility of level three increases, uncertainty about the belief at level two also increases.

The RW model is commonly used for fear conditioning. In this model, beliefs about cue-outcome associations are updated with a fixed learning rate using the PE, where the PE is computed as the difference between the expected and observed outcomes. In our model, p(blame|w) was updated using the following model:

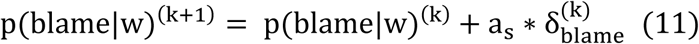

a. When the choice of a participant was wrong, 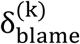 was calculated as

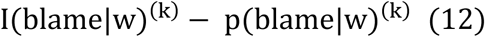
b. When the choice of a participant was correct 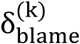 was calculated as

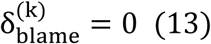

where k is the trial number, a_s_ is the learning rate parameter that ranges from 0 to 1, 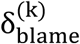 is the blame prediction error (BPE) induced by the difference in expected swearing probability p(blame|w)^(k)^, and I(blame|w)^(k)^ is an indicator function of in the incorrect trial that takes a value of 1 when the feedback is blame and a value of 0 when the feedback is NF. In the correct trial, 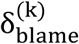 was defined as 0. In this perceptual model, the correct probability 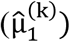 and p(blame|w)^(k)^ were simultaneously updated using information about whether the choice was incorrect and whether the feedback was blame.

The response model employed a modified Softmax function such that the inverse temperature that controls the degrees of change in the trial-by-trial decision noise as functions of uncertainty and p(blame|w) were derived from the HGF3 and RW models, respectively. In this function, the inverse temperature that controls the slope of the function and is related to the degree of decision noise (Wilson et al., 2014) was changed in a trial-by-trial manner as a functions of uncertainty and p(blame|w) as follows:

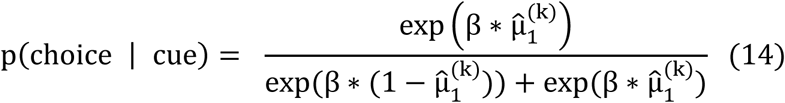

where β is the function of 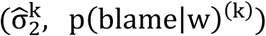 such that

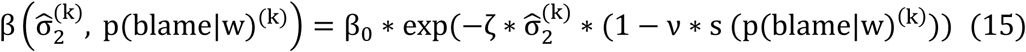

and β_0_ is the baseline inverse temperature, 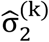 is the estimation of the uncertainty derived from the perceptual model (HGF3) and ζ is the parameter that controls the degree of UDN, which was bounded > 0 by estimating the log space. If the value of this parameter is large, then participants are likely to make suboptimal choices when uncertainty is high. Finally, ν is the parameter that determines the degree of UDN suppression as p(blame|w) increases, which ranges from −1 to 1, as estimated in the inverse hyperbolic tangent space. Using this response function, an increase in uncertainty decreased the slope of the response function, thereby increasing decision noise. Furthermore, p(blame|w) controlled this UDN, which indicates a control of decision noise proportional to the uncertainty level. This strategy enables a flexible and effective control of decision noise rather than always exerting the same level of decision noise control, particularly given the effort required for this control. If ν is positive for certain participants, then those participants suppressed UDN based on p(blame|w), while participants with a negative ν increased UDN.

A detailed description of the priors of the parameters used in the CUDN models are provided in the S5 Text. Moreover, the prior/posterior means and variances of the parameters are listed in S2 Table. Finally, tests of model robustness, including the results of parameter recovery simulation and parameter correlation analysis, are provided in the S6 Text. Briefly, parameters were reliably recovered, and the parameter *v* did not correlate with other parameters, suggesting that this parameter was reliably estimated (S1 Fig). Furthermore, simulated responses recovered a significantly increased choice accuracy in the high-blame blocks (Fig 2D).

In addition to the CUDN model, we considered other models, including 1) a group of models that did not consider the influence of p(blame|w) (RWS, HGF3S and UDN models), 2) a group models in which p(blame|w) has a negative decision value (RWS-N, HGF3S-N, and UDN-N models), and 3) a model in which p(blame|w) functions as a controller of UDN and has a negative decision value (CUDN-N). Detailed descriptions of these models are provided in the S4 Text. Finally, eight models, including the CUDN, RWS, HGF3S, UDN, RWS-N, HGF3S-N, UDN-N and CUDN-N models, were designed. Importantly, in the models where p(blame|w) has a negative value, p(blame|w) consequently decreased the decision noise, regardless of the current uncertainty level (uncertainty-independent control of decision noise). Details of these models are provided in the S4 Text.

### Model fitting and Bayesian model selection

Eight computational models were fit to the participants’ behavior using the TAPAS toolbox. First, the priors of each model were specified (S5 Text, S2 Table), and the Maximum-a-posteriori parameters of each model were obtained by optimizing the log-joint probability function of the prior and the generative model (each computational model) given the response data using the quasi-Newton optimization algorithm (Shanno, 1970). Then, the negative variational free energy, which was an approximation of the log-model evidence was calculated using the Laplace approximation (C. D. Mathys et al., 2014). Using this negative variational free energy as the proxy for the log-model evidence, eight models were compared using random-effects Bayesian model selection (RFX-BMS) to identify the best model that would explain participants’ behaviors, which procedure was implemented using the VBA toolbox (Daunizeau et al., 2014). This RFX-BMS procedure treats the occurrence of the model within each participant as a multinomial random variable with a Dirichlet conjugate prior (Rigoux, Stephan, Friston, & Daunizeau, 2014). By estimating the parameters of the Dirichlet posterior, we were able to calculate the estimated model frequencies and the exceedance probabilities (EPs) of one model being more likely than another (Stephan et al., 2009). Furthermore, by correcting the EP for the Bayesian omnibus risk, which quantifies the probability that the model frequencies are equal, we obtained the protected EPs (PEPs) of the models. These PEPs were calculated using the equations presented in a previous paper (Rigoux et al., 2014). Finally, the model attribution (i.e., the posterior probability of each model for every participant) was calculated (Daunizeau et al., 2014).

### Relationship between UDN suppression and the increase in choice accuracy induced by blame expectation

Because we hypothesized that the suppression of UDN when p(blame|w) is high would help participants make more accurate decisions, we tested whether the participants with a large value for the parameter *v*, indicating a substantial suppression of UDN by p(blame|w), achieved a greater increase in accuracy in the high-blame blocks than participants with small or negative values for *v*. This result was accomplished by performing a correlation analysis between the increase in accuracy and *v*. Furthermore, we expected that the increase in accuracy would exist only in the groups of individuals who suppressed UDN (positive *v*), while participants with a negative *v* would not show increased accuracy in the high-blame blocks. Therefore, we performed separate two-tailed paired *t*-tests of choice accuracy between high- and low-blame blocks in a group with a positive *v* and a group with a negative *v*.

### Model-based fMRI analysis

We performed a model-based fMRI analysis based on the CUDN model to investigate the neural substrate of suppression of UDN under p(blame|w). We performed parametric modulation analyses (N = 41) using the trial-by-trial trajectories of model variables of interest for each subject as parametric modulators of a first-level general linear model (GLM) in Statistical Parametric Mapping 12 (SPM12) (Penny, Friston, Ashburner, Kiebel, & Nichols, 2011). Variables of interest at the onset of the cue included trial-by-trial p(blame|w) and uncertainty that influenced decision noise in the CUDN model. Moreover, we included the trial-by-trial suboptimal value (namely, a smaller value between values (probability) of two options) that is related to a suboptimal choice. Additionally, BPE was added as a parametric modulator. Finally, 24 regressors of no interest, including 6 motion regressors and 18 physiological regressors, were constructed using the TAPAS PhysIo toolbox (Kasper et al., 2017). Then, GLM estimates of the first level were used as an input for the second-level GLM with a one-sample *t*-test design for the regressors of interest using each participant’s sex as a covariate. Additionally, because we did not orthogonalize parametric modulators, especially at the cue, we suspected that multicollinearity might exist between these variables; thus, the beta estimate of p(blame|w) might be unreliable or explain the signal of the other parametric modulators at the cue (uncertainty or suboptimal value). To test whether multicollinearity influenced the estimation of p(blame|w)-related activation, we conducted the same first-level GLM with orthogonalized parametric modulators at the cue, where p(blame|w) was located at the last among the parametric modulators at the cue (order: uncertainty, suboptimal value, and p[blame|w]). This method is similar to that used in the recent paper by Korn and colleagues (Korn & Bach, 2019). Because orthogonalization influences inferences of the beta estimates such that earlier regressors take the shared variances of later regressors (Mumford, Poline, & Poldrack, 2015), we expected that if the activation of p(blame|w) was influenced by multicollinearity, then the activation pattern in the orthogonalized GLM would significantly differ from the original GLM. However, the results of the orthogonal GLM were nearly identical to the original GLM. Thus, the effect of p(blame|w) was less likely to be influenced by multicollinearity. Furthermore, the variance inflation factor (VIF) of p(blame|w) was low (VIF = 1.28), which also shows the low possibility of multicollinearity on p(blame|w)-related activation. Detailed descriptions of the MRI acquisition and fMRI preprocessing steps are provided in the S9 Text and S10 Text.

### Psychophysiological interaction (PPI) analysis

The parametric modulation analysis revealed that the bilateral mPFC was robustly activated by p(blame|w), which is consistent with our hypothesis. Furthermore, we hypothesized that if the mPFC is involved in the suppression of UDN, then the interaction pattern with regions activated by the suboptimal value (S2 Fig and S6 Table) would change as p(blame|w) increases. We performed a PPI analysis (K. Friston et al., 1997) of the mPFC using the generalized psychophysiological interaction (gPPI) toolbox (McLaren et al., 2012) to test this hypothesis. Because the size of the cluster was large and included the mPFCs of both hemispheres, we used both the right/left mPFC ROIs whose center was extracted from the peak voxel of the right/left mPFC within the mPFC cluster. Therefore, two PPI analyses were performed using the right and left mPFCs as seed regions. The mean BOLD signal time courses from the left and right mPFC ROIs of a 6-mm sphere centered at [−8 62 28] and [6 60 32], respectively, were selected from the peak voxels of the mPFC cluster in the p(blame|w) contrast. These mean BOLD signal time series were used as a physiological factor, and p(blame|w) was used as a parametric psychological factor. After identifying clusters whose connectivities were significantly modulated by p(blame|w), we performed a correlation analysis between the mean beta value of the right and left rlPFC clusters obtained from the PPI analysis of the right mPFC with the CUDN model parameter *v* that reflects the degree of suppression of UDN; thus, four correlation analyses were performed. The mean beta values for each cluster were extracted using the MarsBar toolbox (Brett, Anton, Valabregue, & Poline, 2002). The PPI analyses of our study used the CDT applied in the parametric modulation analysis, and no voxels survived. Therefore, we applied the more liberal CDT of p < 0.05, which has been used in previous PPI analyses *(Guediche, Reilly, Santiago, Laurent, & Blumstein, 2016; Outhred et al., 2015; Paret et al., 2016)* and a minimum size of 500 voxels.

### Dynamic causal modeling analysis

According to the results of the parametric modulation analysis and PPI analysis, negative modulation of functional connectivity between the right mPFC and right rlPFC, which was activated by the suboptimal value, had a correlation with *v* while other PPI results did not showed this relationship. While the results of the PPI analyses provided an important clue that the modulation of the neural dynamics between the right mPFC and rlPFC are related to UDN suppression and other regions are not likely to be involved in this process, the PPI analyses only shows only a change of correlational pattern and did not provide possible mechanistic neural models associated with the suppression of UDN. Furthermore, the correlation with the parameter *v* was not very strong given that we performed four correlation analyses between PPI cluster and *v* (p = 0.029, which was not significant after applying the Bonferroni correction). We expected that investigating more specific neural dynamics model of the regions associated with blame expectation using the DCM and determining the components relevant to the suppression of UDN within that neural dynamics model would help us to clarify the neural mechanism of UDN suppression. Therefore, we performed a DCM analysis (K. J. Friston et al., 2003). In DCM analyses using SPM12 software, it is typical to choose ROIs among the regions showing an experimental effect (Zeidman et al., 2019); thus, we chose two ROIs based on our experimental effects. Specifically, the right mPFC was activated based on p(blame|w), and the right rlPFC showed a negative PPI effect with the right mPFC based on p(blame|w). Furthermore, the ROIs were 8-mm spheres centered on [6 60 32] and [28 66 6], respectively, and were the peak voxels detected in the p(blame|w) contrast and PPI analysis. We extracted the principal eigenvariate from each participant within the fixed-centered ROI where the voxels used in these steps were created using a liberal threshold of an uncorrected p-value of < 0.1 based on the first-level GLM results, which is the same threshold used in a previous study (Legon et al., 2016), and adjusted for the effect of interest. However, after this extraction, no voxels survived in nine participants: 3 participants showed no parametric modulation of p(blame|w) in the right mPFC ROI, and 6 participants showed no negative PPI effect in the right rlPFC ROI. Thus, we excluded these participants from the subsequent DCM analysis, and the data from 32 participants were used in the DCM analysis. We tested 16 models that varied in 1) the existence of a fixed connection between two ROIs, assuming that each ROI has a recurrent connection from itself; 2) whether modulatory inputs of the p(blame|w) to each directional connectivity existed; and 3) whether a driving input of p(blame|w) modulated the mPFC (we assumed that p(blame|w) did not act as a driving input to the rlPFC). Graphical descriptions of all models are provided in S5 Fig. Models were compared using RFX-BMS implemented in SPM12 (Penny et al., 2011).

## Data and code availability

Source codes for computational models of behavior are accessible at the 10.6084/m9.figshare.8157896. Other data from the current study are available from the corresponding author upon reasonable request.

## Acknowledgments

This research was supported by the Brain Research Program through the National Research Foundation of Korea (NRF) funded by the Ministry of Science & ICT (NRF - 2016M3C7A1914448 and NRF - 2017M3C7A1031331), Institute for Information & communication Technology Planning & evaluation (IITP - 20170007800031001) and the BK21 Plus Fund (22A20151313464). The funders had no role in study design, data collection and analysis, decision to publish, or preparation of the manuscript. The language of this manuscript was edited by American Journal Experts (AJE). The authors wish to acknowledge Seokho Yoon, Minseob Kim, Dohyun Kim and Geumsuk Shim for performing the psychiatric evaluations of the participants; Haeorm Park, Minseob Kim, Dohyun Kim, Mincheol Kim and Dongwoo Shin for acquiring the MRI data; and Mincheol Kim and Kwangsun Yoo for participating in helpful discussions.

## Supplementary materials

**S1 Table.** Post-experimental survey regarding the influence of expecting blame on participant behavior and mood.

| **Survey Items** | **Number of Participants** |
| --- | --- |
| **Influence of blame expectation on choice** |  |
| 1. Did not think about how blame feedback would be likely to be given by making an incorrect choice. | 3/33 (9.1%) |
| 2. I expected to receive blame feedback when it was likely; however, it did not affect my behavior. | 6/33 (18.2%) |
| 3. When blame feedback was likely after an incorrect choice, I paid more attention when deciding. | 12/33 (36.4%) |
| 4. When blame feedback was likely after an incorrect choice, it was difficult to pay attention to the choice. | 3/33 (9.1%) |
| 5. When blame feedback was likely after an incorrect choice, I made a more deliberate decision (tried to choose an option that would be more likely to be correct). | 13/33 (39.4%) |
| 6. When blame feedback was likely after an incorrect choice, it was difficult to make a decision. | 4/33 (12.1%) |
| 7. When blame feedback was likely after an incorrect choice, I felt like making a riskier choice (chose an option that was likely to be incorrect). | 3/33 (9.1%) |
| **Influence of blame expectation on mood** |  |
| 8. When blame feedback was likely after an incorrect choice, I felt uncomfortable. | 18/33 (54.6%) |
| **Summary of the influence of blame expectation on choice behavior (among 22 participants who reported that their behavior was influenced)** | **Number of Participants** |
| Positively influenced (enhanced attention/facilitated deliberative choice) | 14/22 (63.6%) |
| Negatively influenced (decreased attention/facilitated risky choice/made decisions difficult) | 4/22 (18.2%) |
| Mixed (checked both positive and negative influences) | 4/22 (18.2%) |

**S2 Table.** Description of the parameters of the CUDN model.

| Parameter | Description | Prior mean | Prior SD | Posterior mean | Posterior SD |
| --- | --- | --- | --- | --- | --- |
| $\omega$ | Tonic volatility parameter | -3 | 4 | -2.04 | 0.90 |
| $\log(\vartheta)$<br>(fixed) | Meta volatility parameter | -6 | 0 | -6 | 0 |
| $\kappa$ (fixed) | Influence of 3<sup>rd</sup> level on 2<sup>nd</sup> level | 1 | 0 | 1 | 0 |
| $\log(\zeta)$ | UDN parameter | 0 | 1 | 0.15 | 0.49 |
| $\log(\beta_0)$ | Baseline inverse temperature | $\log 48$ | 1 | 3.28 | 0.61 |
| $\text{logit}(a_s)$ | Learning rate of $p(\text{blame}\vert \text{w})$ | 0 | 0.7 | 0.02 | 0.07 |
| * $\tanh^{-1}(\mathbf{v})$ | Suppression of UDN based on $p(\text{blame}\vert \text{w})$ | 0 | 0.7 | 0.05 | 0.33 |

\* $\tanh^{-1}$ : inverse hyperbolic tangent function

**S3 Table.** Results of the Poisson regression analysis using parameters of the CUDN model explaining PHQ-9 and GAD-7 outcomes.

| Model parameter | Coefficient ($\beta$) of the PHQ-9 model | Coefficient ($\beta$) of the GAD-7 model |
| --- | --- | --- |
| $\omega$ | 0.25 (t (37) = 2.83, p = 0.005) | 0.27 (t (37) = 2.45, p = 0.014) |
| $\zeta$ | 0.21 (t (37) = 3.03, p = 0.002) | 0.09 (t (37) = 0.90, p = 0.367) |
| $\beta_0$ | -0.13 (t (37) = -1.74, p = 0.082) | -0.27 (t (37) = -2.59, p = 0.01) |
| $a_s$ | -0.11 (t (37) = -1.21, p = 0.226) | -0.73 (t (37) = -4.09, p < 0.001) |
| $\mathbf{v}$ | <u>-0.248 (t (37) = -2.88, p = 0.004)</u> | <u>-0.589 (t (37) = -3.92, p < 0.001)</u> |
| Sex | -0.70 (t (37) = -3.63, p < 0.001) | -1.05 (t (37) = 4.06, p < 0.001) |

**S4 Table.** Model attribution for each participant.

|  | Model attribution (probability) |  |  |  |  |  |  |  |
| --- | --- | --- | --- | --- | --- | --- | --- | --- |
|  | RW | HGF3S | UDN | RWS-N | HGF3S-N | UDN-N | CUDN-N | CUDN |
| **S1** | 0.00 | 0.00 | 0.12 | 0.00 | 0.00 | 0.01 | 0.00 | 0.88 |
| **S2** | 0.00 | 0.00 | 0.11 | 0.00 | 0.00 | 0.17 | 0.00 | 0.72 |
| **S3** | 0.00 | 0.00 | 0.07 | 0.00 | 0.00 | 0.43 | 0.00 | 0.50 |
| **S4** | 0.00 | 0.00 | 0.02 | 0.00 | 0.00 | 0.88 | 0.00 | 0.10 |
| **S5** | 0.00 | 0.00 | 0.13 | 0.00 | 0.00 | 0.00 | 0.00 | 0.86 |
| **S6** | 0.00 | 0.00 | 0.14 | 0.00 | 0.00 | 0.01 | 0.00 | 0.85 |
| **S7** | 0.00 | 0.00 | 0.04 | 0.00 | 0.00 | 0.08 | 0.00 | 0.88 |
| **S8** | 0.00 | 0.00 | 0.11 | 0.00 | 0.00 | 0.27 | 0.00 | 0.62 |
| **S9** | 0.00 | 0.00 | 0.01 | 0.00 | 0.00 | 0.86 | 0.00 | 0.13 |
| **S10** | 0.00 | 0.00 | 0.03 | 0.00 | 0.00 | 0.77 | 0.00 | 0.20 |
| **S11** | 0.00 | 0.00 | 0.13 | 0.00 | 0.00 | 0.06 | 0.00 | 0.81 |
| **S12** | 0.00 | 0.00 | 0.12 | 0.00 | 0.00 | 0.02 | 0.00 | 0.85 |
| **S13** | 0.00 | 0.00 | 0.13 | 0.00 | 0.00 | 0.00 | 0.00 | 0.86 |
| **S14** | 0.00 | 0.00 | 0.13 | 0.00 | 0.00 | 0.00 | 0.00 | 0.87 |
| **S15** | 0.00 | 0.00 | 0.13 | 0.00 | 0.00 | 0.02 | 0.00 | 0.85 |
| **S16** | 0.00 | 0.00 | 0.12 | 0.00 | 0.00 | 0.09 | 0.00 | 0.79 |
| **S17** | 0.00 | 0.00 | 0.14 | 0.00 | 0.00 | 0.03 | 0.00 | 0.83 |
| **S18** | 0.00 | 0.00 | 0.12 | 0.00 | 0.00 | 0.00 | 0.00 | 0.88 |
| **S19** | 0.00 | 0.00 | 0.11 | 0.00 | 0.00 | 0.20 | 0.00 | 0.69 |
| **S20** | 0.00 | 0.00 | 0.06 | 0.00 | 0.00 | 0.02 | 0.00 | 0.91 |
| **S21** | 0.00 | 0.00 | 0.06 | 0.00 | 0.00 | 0.52 | 0.00 | 0.42 |
| **S22** | 0.00 | 0.00 | 0.09 | 0.00 | 0.00 | 0.29 | 0.00 | 0.62 |
| **S23** | 0.00 | 0.00 | 0.07 | 0.00 | 0.00 | 0.18 | 0.00 | 0.74 |
| **S24** | 0.00 | 0.00 | 0.08 | 0.00 | 0.00 | 0.05 | 0.00 | 0.88 |
| **S25** | 0.00 | 0.00 | 0.09 | 0.00 | 0.00 | 0.29 | 0.00 | 0.61 |
| **S26** | 0.00 | 0.00 | 0.00 | 0.00 | 0.00 | 1.00 | 0.00 | 0.00 |
| **S27** | 0.00 | 0.00 | 0.13 | 0.00 | 0.00 | 0.00 | 0.00 | 0.87 |
| **S28** | 0.00 | 0.00 | 0.05 | 0.00 | 0.00 | 0.62 | 0.00 | 0.33 |
| **S29** | 0.00 | 0.00 | 0.02 | 0.00 | 0.00 | 0.01 | 0.00 | 0.97 |
| **S30** | 0.00 | 0.00 | 0.06 | 0.00 | 0.00 | 0.51 | 0.00 | 0.43 |
| **S31** | 0.00 | 0.00 | 0.08 | 0.00 | 0.00 | 0.41 | 0.00 | 0.52 |
| **S32** | 0.00 | 0.00 | 0.12 | 0.00 | 0.00 | 0.02 | 0.00 | 0.85 |
| **S33** | 0.00 | 0.00 | 0.12 | 0.00 | 0.00 | 0.04 | 0.00 | 0.84 |
| **S34** | 0.00 | 0.00 | 0.12 | 0.00 | 0.00 | 0.01 | 0.00 | 0.87 |
| **S35** | 0.00 | 0.00 | 0.00 | 0.00 | 0.00 | 0.97 | 0.00 | 0.03 |
| **S36** | 0.00 | 0.00 | 0.11 | 0.00 | 0.00 | 0.19 | 0.00 | 0.70 |
| **S37** | 0.00 | 0.00 | 0.13 | 0.00 | 0.00 | 0.00 | 0.00 | 0.86 |
| **S38** | 0.00 | 0.00 | 0.07 | 0.00 | 0.00 | 0.49 | 0.00 | 0.44 |
| **S39** | 0.00 | 0.00 | 0.14 | 0.00 | 0.00 | 0.00 | 0.00 | 0.86 |
| **S40** | 0.00 | 0.00 | 0.11 | 0.00 | 0.00 | 0.19 | 0.00 | 0.70 |
| **S41** | 0.00 | 0.00 | 0.13 | 0.00 | 0.00 | 0.00 | 0.00 | 0.87 |
| **S42** | 0.00 | 0.00 | 0.12 | 0.00 | 0.00 | 0.05 | 0.00 | 0.82 |
| **S43** | 0.00 | 0.00 | 0.10 | 0.00 | 0.00 | 0.19 | 0.00 | 0.71 |
| **S44** | 0.00 | 0.00 | 0.14 | 0.00 | 0.00 | 0.00 | 0.00 | 0.86 |
| **S45** | 0.00 | 0.00 | 0.00 | 0.00 | 0.00 | 0.99 | 0.00 | 0.01 |
| **S46** | 0.00 | 0.00 | 0.11 | 0.00 | 0.00 | 0.01 | 0.00 | 0.88 |

\*S1-S46: Participants 1 to 46

**S5 Table.** Regions activated by BPE.

| **Cluster** | **Cluster**<br>**p-value**<br>**(FWE**<br>**corrected**<br>**)** | **No. of**<br>**voxels** | **Region name**<br>**(AAL)** | **MNI**<br>**coordinates**<br>**(x, y, z)** | **Voxel**<br>**Z-Value**<br>**(uncorrected**<br>**p-value)** |
| --- | --- | --- | --- | --- | --- |
| **ACC/MCC** | < 0.001 | 1878 | Cingulate_Mid_L | -6, -20, 32 | 5.45<br>(< 0.001) |
|  |  |  | Cingulate_Ant_L | -4, 26, 26 | 5.31<br>(< 0.001) |
| **SMA** | < 0.001 | 1272 | Supp_Motor_Area_L | -4, 10, 62 | 5.89<br>(< 0.001) |
| **PAG/cerebellum/**<br>**thalamus** | < 0.001 | 310 | PAG | 4, -30, -6 | 4.32<br>(< 0.001) |
|  |  |  | Vermis_3 | 4, -32, -4 | 5.65<br>(< 0.001) |
|  |  |  | Thalamus_L | -4, -28, 0 | 4.88<br>(< 0.001) |
| **Right**<br>**MTG/insula** | < 0.001 | 5622 | Temporal_Pole_Mid_R | 50, 20, -30 | 6.58<br>(< 0.001) |
|  |  |  | Insula_R | 28, 16, -20 | 6.26<br>(< 0.001) |
| **Left IFG** | < 0.001 | 7900 | Frontal_Inf_Tri_L | -52, 28, 4 | 7.39<br>(< 0.001) |
| **Right fusiform**<br>**gyrus** | < 0.001 | 1812 | Fusiform_R | 44, -60, -18 | 6.58<br>(< 0.001) |
| **Left fusiform**<br>**gyrus** | < 0.001 | 2089 | Fusiform_L | -42, -54, -24 | 6.39<br>(< 0.001) |
| **Right precentral**<br>**gyrus** | < 0.001 | 420 | Precentral_R | 44, 2, 44 | 5.39<br>(< 0.001) |
| **Left precentral**<br>**gyrus** | < 0.001 | 648 | Precentral_L | -40, 2, 46 | 5.09<br>(< 0.001) |
| **Left precuneus** | < 0.001 | 2080 | Precuneus_L | -8, -50, 38 | 5.33<br>(< 0.001) |
| **mPFC** | < 0.001 | 1511 | Frontal_Sup_Medial_L | -4, 52, 28 | 4.88<br>(< 0.001) |
| **Right**<br>**hippocampus** | < 0.001 | 131 | Hippocampus_R | 8, -12, -12 | 4.81<br>(< 0.001) |

**S6 Table.** Regions activated by the suboptimal value.

| **Cluster** | **Cluster**<br>**p-value**<br>**(FWE**<br>**corrected)** | **No. of**<br>**voxels** | **Region name**<br>**(AAL)** | **MNI**<br>**coordinates**<br>**(x, y, z)** | **Voxel**<br>**Z-Value**<br>**(uncorrected**<br>**p-value)** |
| --- | --- | --- | --- | --- | --- |
| **Right IFG**<br>**/dmPFC** | < 0.001 | 8069 | Frontal_Inf_Orb_2_R | 34, 24, -8 | 7.20<br>(< 0.001) |
|  |  |  | Frontal_Sup_Medial_R | 6, 32, 40 | 6.48<br>(< 0.001) |
| **Right rIPFC** | < 0.001 | 1840 | Frontal_Mid_2_R | 36, 62, -8 | 5.09<br>(< 0.001) |
| **Left rIPFC** | < 0.001 | 440 | Frontal_Mid_2_L | -30, 56, 8 | 4.75<br>(< 0.001) |
| **Left rIPFC** | 0.05 | 123 | OFCant_L | -18, 58, -14 | 4.19<br>(< 0.001) |
| **right IPL** | < 0.001 | 2237 | Parietal_Inf_R | 38, -46, 42 | 5.54<br>(< 0.001) |
| **Left IPL** | < 0.001 | 1470 | Parietal_Inf_L | -32, -50, 40 | 6.05<br>(< 0.001) |
| **Left insula** | < 0.001 | 910 | Insula_L | -28, 22, -6 | 5.93<br>(< 0.001) |
| **Thalamus/caudate** | < 0.001 | 1931 | Thalamus_R | 14, -12, 8 | 5.48<br>(< 0.001) |
|  |  |  | Caudate_R | 16, -2, 16 | 5.34<br>(< 0.001) |
| **Precuneus** | < 0.001 | 566 | Precuneus_R | 6, -70, 44 | 5.32<br>(< 0.001) |
| **mPFC** | < 0.001 | 1511 | Frontal_Sup_Medial_L | -4, 52, 28 | 4.88<br>(< 0.001) |
| **Right hippocampus** | < 0.001 | 131 | Hippocampus_R | 8, -12, -12 | 4.81<br>(< 0.001) |
|  |  |  | Temporal_Mid_R | 50, -18, -16 | 4.94<br>(< 0.001) |
| **Left precentral gyrus** | < 0.001 | 656 | Precentral_L | -52, 12, 38 | 4.81<br>(< 0.001) |
| **Left MFG** | 0.006 | 193 | Frontal_Mid_2_L | -28, -2, 50 | 4.00<br>(< 0.001) |
| **Cerebellum** | < 0.001 | 711 | Cerebellum_Crus1_R | 40, -70, -28 | 5.39<br>(< 0.001) |
| **Cerebellum** | < 0.001 | 2824 | Cerebellum_Crus1_L | -32, -60, -32 | 5.51<br>(< 0.001) |
| **Cerebellum** | < 0.001 | 225 | Vermis_3 | 2, -44, -22 | 3.98<br>(< 0.001) |

**S7 Table.** Regions activated by the optimal value. (maximum value between two options)

| **Cluster** | **Cluster p-value (FWE corrected)** | **No. of voxels** | **Region name (AAL)** | **MNI coordinates (x, y, z)** | **Voxel Z-Value (uncorrected p-value)** |
| --- | --- | --- | --- | --- | --- |
| **mOFC** | < 0.001 | 2298 | Frontal_Inf_Orb_2_R | 34, 24, -8 | 7.197<br>($< 0.001$) |
| **Precuneus** | < 0.001 | 9147 | Precuneus_L | -6, -56, 16 | 5.88<br>($< 0.001$) |
| **Right hippocampus** | < 0.001 | 150 | Hippocampus_R | 28, -16, -20 | 5.58<br>($< 0.001$) |
| **Left insula** | < 0.001 | 2237 | Insula_L | -36, -28, 24 | 5.25<br>($< 0.001$) |
| **Left dlPFC** | < 0.001 | 541 | Frontal_Sup_2_L | -20, 32, 42 | 4.86<br>($< 0.001$) |
| **Left insula** | < 0.001 | 910 | Insula_L | -28, 22, -6 | 5.93<br>($< 0.001$)<sup>1</sup> |
| **Left vmPFC** | 0.005 | 201 | Frontal_Sup_Medial_L | -12, 62, 8 | 4.40<br>($< 0.001$) |
| **Right cuneus** | 0.021 | 152 | Cuneus_R | 14, -92, 22 | 4.18<br>($< 0.001$) |

**S8 Table.** PPI analysis results of the bilateral mPFC ROI using p(blame|w) as a psychological factor.

| Cluster | Cluster<br>p-value<br>(FWE<br>corrected) | No. of<br>voxels | Region name<br>(AAL) | MNI<br>coordinates<br>(x, y, z) | Voxel<br>Z-Value<br>(uncorrected<br>p-value) |
| --- | --- | --- | --- | --- | --- |
| Negative PPI with right mPFC seed |  |  |  |  |  |
| Right<br>rIPFC | 0.039 | 1320 | Frontal_Sup_2_R | 28, 66, 6 | 3.47<br>($< 0.001$) |
| Cuneus<br>/Rolandic<br>operculum | $< 0.001$ | 5498 | Rolandic_Oper_R | 34, -42, 22 | 3.76<br>($< 0.001$) |
| | | | Cuneus_R | 4, -80, 30 | 3.70<br>($< 0.001$) |
| Positive PPI with left mPFC seed |  |  |  |  |  |
| Left SMA | $< 0.001$ | 2854 | Supp_Motor_Area_<br>L | -6, 20, 64 | 3.60<br>($< 0.001$) |
| Negative PPI with left mPFC seed |  |  |  |  |  |
| Right<br>cuneus | 0.005 | 1863 | Cuneus_R | 12, -82, 44 | 3.45<br>($< 0.001$) |

**S9 Table.** PPI results of the PAG ROI using BPE as a psychological factor.

| Cluster | Cluster p-value (FWE corrected) | No. of voxels | Region name (AAL) | MNI coordinates (x, y, z) | Voxel Z-Value (uncorrected p-value) |
| --- | --- | --- | --- | --- | --- |
| **aMCC** | 0.046 | 1747 | Cingulate_Mid_R | 8, 18, 40 | 3.32<br>($< 0.001$) |
|  |  |  | Cingulate_Ant_L | -6, 12, 28 | 2.91<br>(0.001) |

**S1 Fig.**
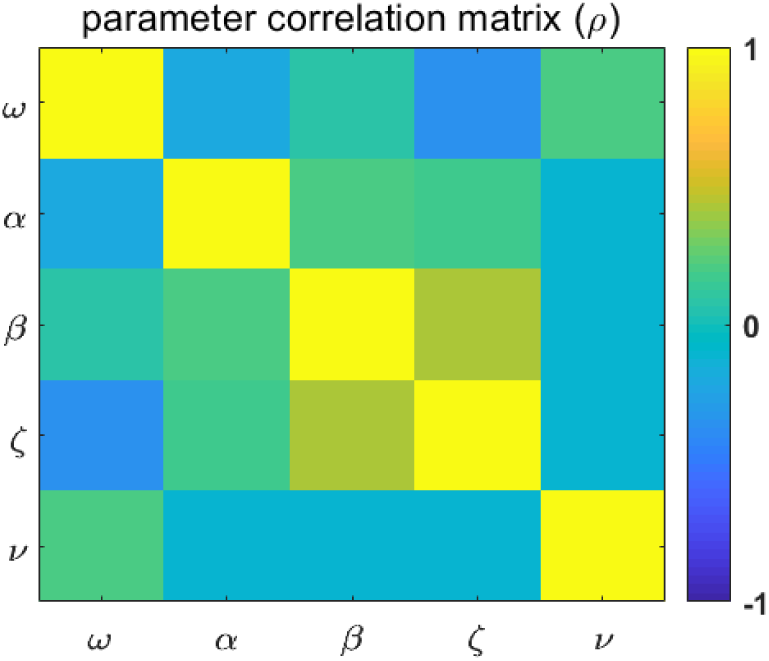
Parameter correlation matrix of the CUDN model. Two significant correlations were identified between these parameters, including a correlation between *ζ* and *β*_0_ (p = 0.004) as well as between *ζ* and ω (p = 0.018). Notably, no parameters were correlated with the parameter of interest, *v*.

**S2 Fig.**
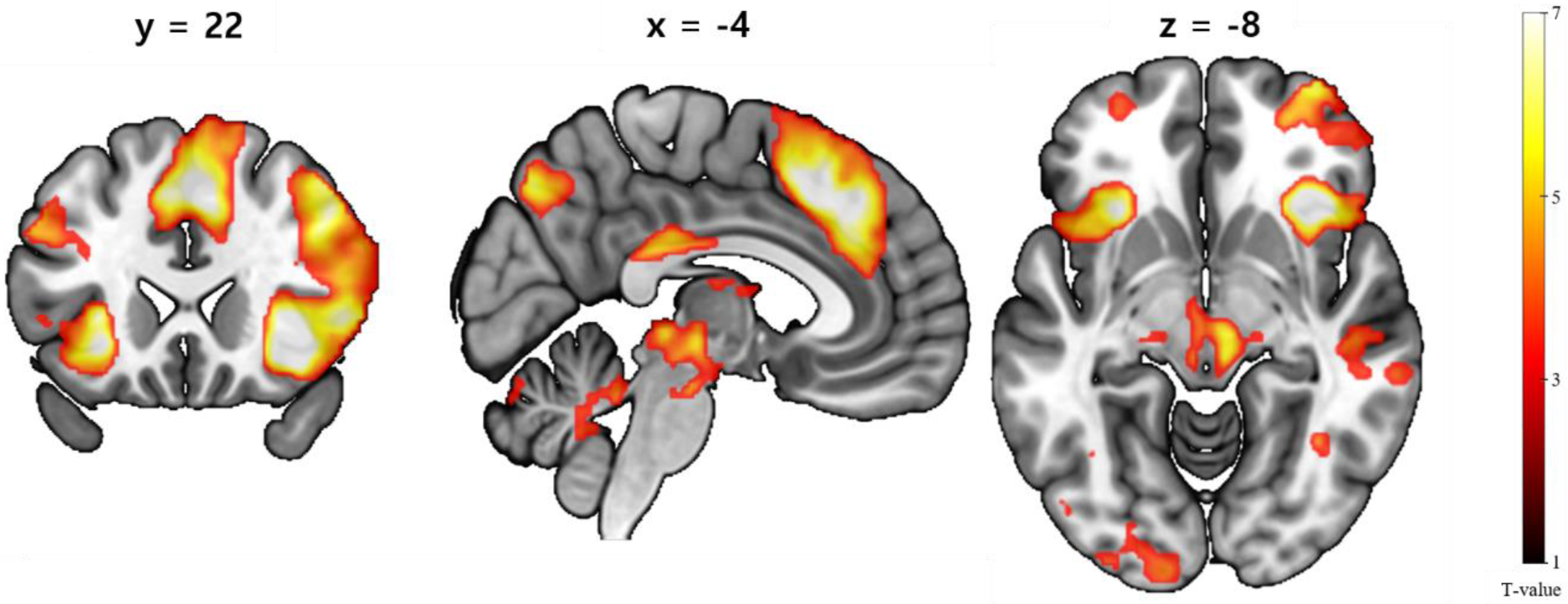
Regions activated by the suboptimal value. Bilateral rlPFC clusters, including the posterior portion of the mPFC including dACC, and bilateral IPL regions were activated by the suboptimal value (S6 Table).

**S3 Fig.**
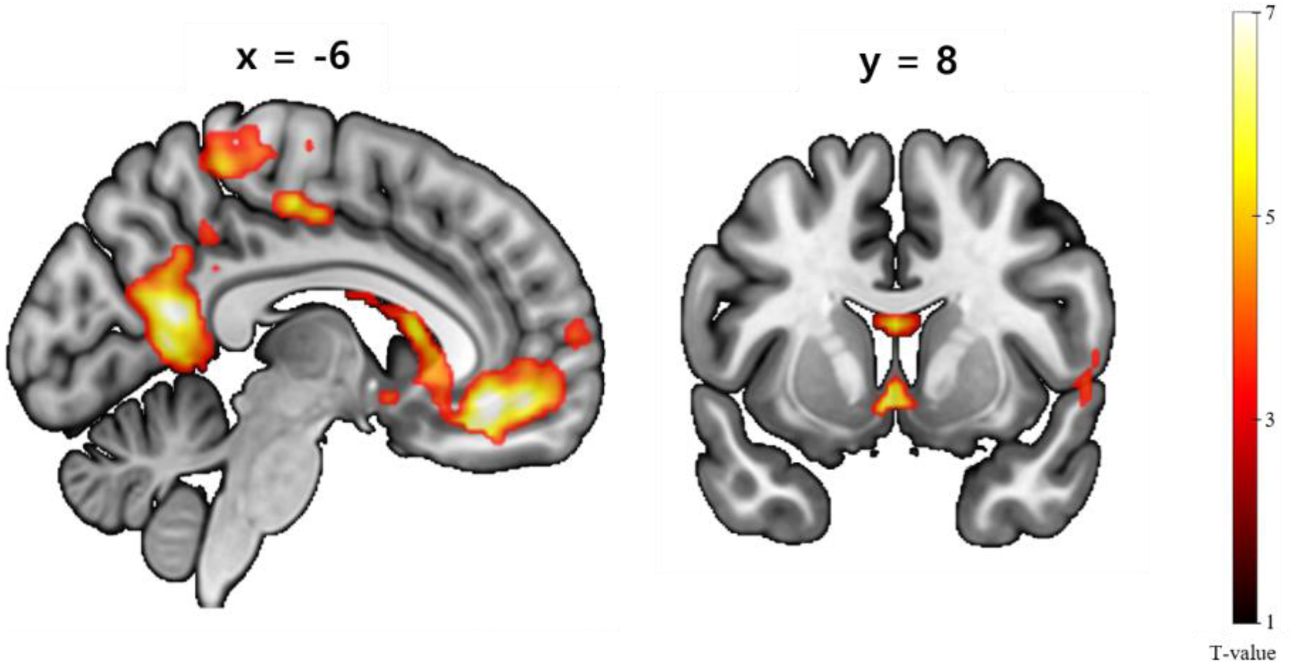
Regions activated by the optimal value. Medial OFC, precuneus, hippocampus and left insula was activated by an optimal value, which is the maximum value between two options (max(p(correct|cue1), p(correct|cue2)) S7 Table).

**S4 Fig.**
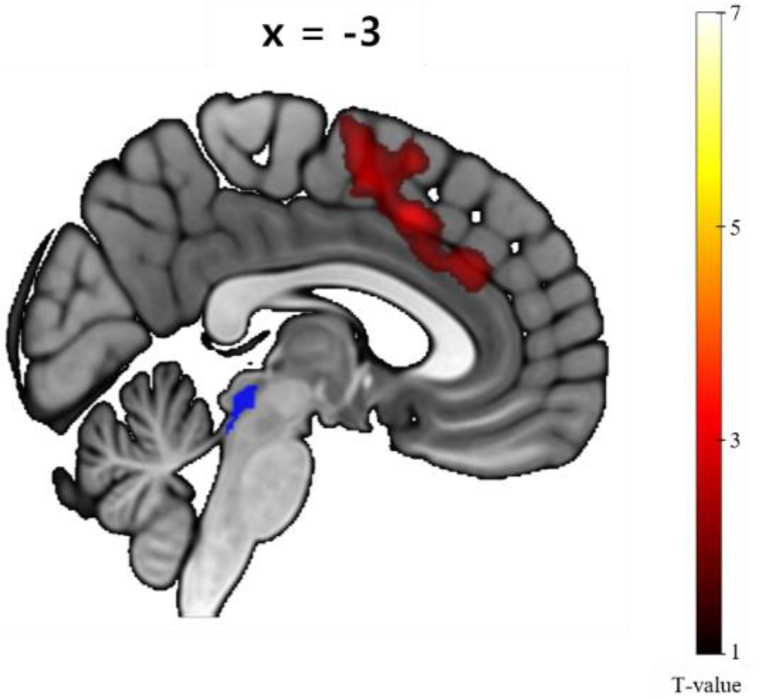
PAG ROI (blue) and the regions showing positive PPI with PAG. BPE positively modulated the functional connectivity between the PAG and aMCC (S9 Table).

**S5 Fig.**
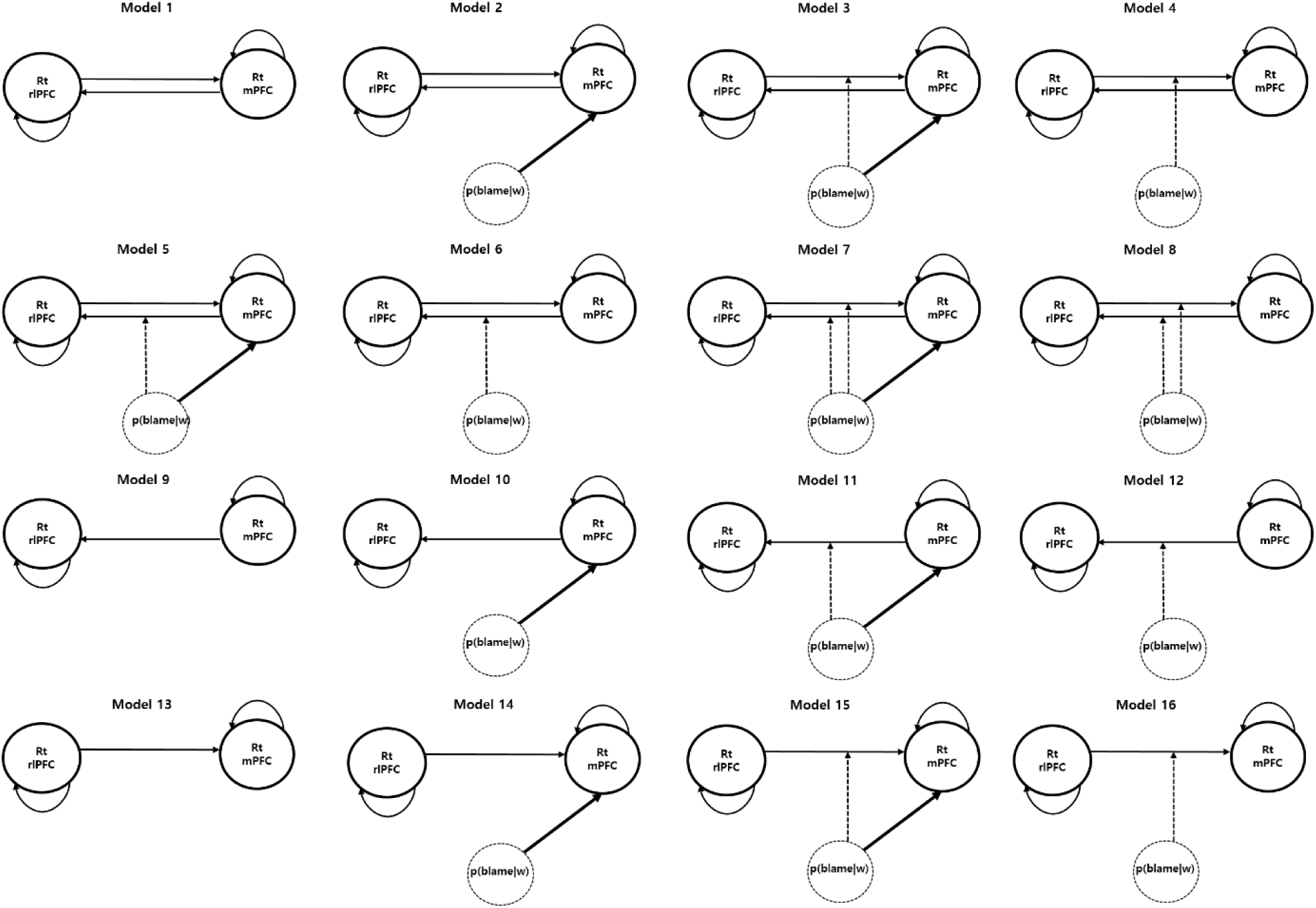
All DCM models used in the Bayesian model selection. We tested 16 DCM models that varied in 1) the existence of a fixed connection between two ROIs, assuming that each ROI has a recurrent connection from itself; 2) whether modulatory inputs of the p(blame|w) to each directional connectivity existed; and 3) whether a driving input of p(blame|w) modulated the mPFC (we assumed that p(blame|w) did not act as a driving input to the rlPFC). Bayesian model selection result showed that model 7 was the winning model. This model showed bidirectional fixed connections between the right mPFC and rlPFC which connections were modulated by p(blame|w) and p(blame|w) also acted as a driving input to the mPFC (Fig 5).

## Supporting Text

### Supporting Text 1. Stimuli and post-experimental survey

We used one neutral and one angry facial expression from the same person for NF and blame from the Korean Facial Expressions of Emotions (KOFEE) database (Park et al., 2011). Four swear words that were validated and used in a previous study (Lee et al., 2017) were also employed. An angry face and the four swear words were combined to make four strong blame stimuli. To verify the various effects of these new stimuli on the tasks, including any effect on decision making and mood, we asked participants to complete a post-experimental survey.

### Supporting Text 2. Post-experimental survey results

In the first section of the post-experimental survey, participants were asked to rate the valence of the secondary feedback stimuli composed of combined faces and words, including a neutral face with the word “Correct”, a neutral face with the word “Wrong” and an angry face with the four swear words. Therefore, 45 participants rated six face-word pairs (1 participant did not complete the first part of the post-experimental survey) using a seven-point Likert scale ranging from −3 to +3. The mean valence of the neutral face with the word “Correct” was 0.38 (± 0.75); that of the neutral face with the word “Wrong” (NF feedback) was − 0.47 (± 0.69); and those of the angry face with four swear words (blame) were −1.96 (± 0.77), −1.84 (± 0.82), −2.13 (± 0.81), and −2.47 (± 0.75). Moreover, the valence of blame was significantly lower than that of NF (two-tailed paired *t*-test, t(44) = −16.7, p < 0.001, CIs: −1.83 to −1.43, *d* = −2.5), and the valence of NF was significantly lower than that of correct feedback (two-tailed paired *t-* test, t(44) = −5.95, p < 0.001, CIs: −1.13 to −0.56, *d* = −0.9).

The second section of the survey asked participants about the influence of expecting blame after making an incorrect choice on their choice behavior and mood. A total of 36 participants completed this survey, and the responses of 3 participants were excluded because they were unreliable (e.g., YAD_10050 reported that he did not think about how blame feedback is likely to be given by making an incorrect choice but he also reported that when he thought that he was highly likely to receive a blame due to wrong choice, he felt like he was making a riskier choice, which is a contradictory answer). Thus, 33 participants’ data were used for this analysis. Of the 33 participants, only 3 reported that they did not consciously think about blame feedback in the event of an incorrect choice; 6 reported that the expectation of blame after an incorrect choice did not affect them; and 2 reported that it affected their mood but not their choice. Of the 22 participants who reported that the expectation of blame after making an incorrect choice influenced their choice behavior, 14 were influenced in a positive way, and 4 reported that they were negatively influenced. The full survey items are listed in S1 Table.

### Supporting Text 3. Behavioral analysis of the influence of task design on behavior

In addition to testing the accuracy difference between the high- and low-blame blocks, we tested whether the design of our task influenced participants’ behavior as we intended using a mixed-effect logistic regression analysis. First, we tested whether participants’ responses were influenced by the designed probability structure of p(correct|cue), which frequently changed during task and therefore might be difficult to follow. To test this possibility, we performed a mixed-effect logistic regression analysis with participant response as the dependent variable, the designed probability structure of p(correct|cue) as the fixed effect, and participant and sex as random effects. The results showed that designed probability structure significantly influenced participant response (beta = 3.27, t(10907) = 36.67, p < 0.001).

Second, we tested whether our task design influenced participants’ suboptimal choice without using explicit computational model of the behavior before performing the model-driven behavioral analysis. In particular, we were interested in whether the uncertainty-related design and designed blame probability (by making an incorrect choice) influenced the choice of the “bad” option, which was designed to have lower probability of being correct. According to our hypothesis, we expected that the uncertainty-related task design would influence participants to make a bad choice and that designed blame probability would decrease this behavioral tendency. Note that this bad choice differs from an actual suboptimal choice of an agent, which was defined as the choice of a cue with a lower probability of being correct. Here, “probability” is the subjective belief or inference of an agent that cannot be observed directly but can be estimated using a computational model. In the task, the frequent change or reversal of p(correct|cue) (e.g., from 0.2 to 0.8) was designed to induce an uncertainty with regard to the inferred p(correct|cue) of participants. We expected that the uncertainty induced by reversal would be reduced until the next reversal because the agent would learn about the current (designed) p(correct|cue) (De Berker et al., 2016). Therefore, we defined the “reversal effect” as a variable that has a maximum value of 1 at the trial after the reversal and decreases in an inversely proportional manner with regard to the number of trials from the current reversal until the next reversal. We used the reversal effect as a proxy for “designed” uncertainty. We performed a mixed-effect logistic regression analysis using the choice of a bad option as the dependent variable. The fixed-effect variables were the reversal effect, designed blame probability, the difference of designed p(correct|cue1) and p(correct|cue2) (we called this variable “designed DV”). Finally, participant and sex were used as random effects.

As we expected, the reversal effect significantly increased the choice of the bad option (beta = 0.73, t[10902] = 25.98, p < 0.001) because participants have less information about the reversed probability immediately after the reversal and learn about it gradually. However, the reversal effect showed a significant negative interaction with designed blame probability (beta = −0.13, t[10902] = 4.62, p < 0.001), showing that the reversal effect on the choice of the bad option decreases as the designed blame probability increases, consistent with our expectations.

Importantly, these analysis results provide only a clue that uncertainty increases the likelihoods of a suboptimal choice and that the expectation of being blamed reduces uncertainty-induced suboptimal choice because the variables used in this analysis were related to only the task design and did not directly reflect participants’ internal computation of the decision variables (e.g., uncertainty) or suboptimal or optimal decisions based on these variables. Therefore, in the next step we performed a model-based behavior analysis based on the trial-by-trial trajectories of the decision variables estimated by fitting adequate computational models to the response data.

### Supporting Text 4. Description of other behavioral models, excluding the CUDN model

In addition to the CUDN model, we considered other models including 1) a group of models that did not consider the influence of p(blame|w) (RWS, HGF3S and UDN models) and 2) a group models in which p(blame|w) has a negative value (RWS-N, HGF3S-N, UDN-N, and CUDN-N models). The following group of models that did not consider the influence of p(blame|w) included

1. RW with a basic Softmax model (RWS) such that the perceptual model calculated p(correct|cue) using the RW rule, and the response model is the basic Softmax model
2. HGF3 with a basic Softmax model (HGF3S) such that the perceptual model calculated p(correct|cue) using the HGF3 model, and the response model is the basic Softmax model
3. HGF3 with an uncertainty-induced decision noise model (UDN model) such that the perceptual model calculated p(correct|cue) using the HGF3 model, and the response model is the modified Softmax model in which uncertainty increases decision noise.

The basic Softmax model was calculated as follows:

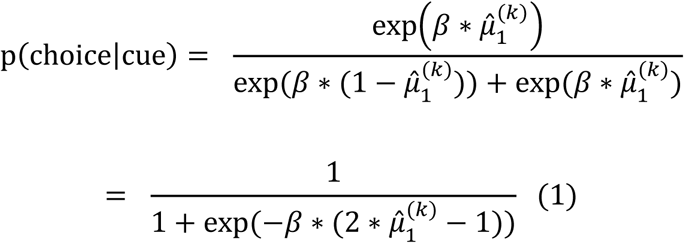

where 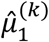 indicates p(correct|cue) both in the HGF3 model and the RW model, and the inverse temperature *β* is a parameter but not a function because the CUDN model is fixed across trials; thus, it is neither influenced by uncertainty nor p(blame|w). The UDN model used HGF3 as a perceptual model and the modified Softmax model as a response function, where the inverse temperature is the function of uncertainty:

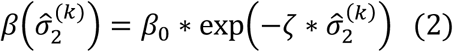

where an uncertainty 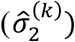 increases decision noise, which similar to the CUDN model, except for the lack of the influence of p(blame|w).

We next constructed a group of models in which p(blame|w) had a negative value. In these models, p(blame|w) was calculated using the same method as in the CUDN model (RW rule), and the total decision value of the cue at trial k (*V*(*cue*)^(*k*)^) for the decision was calculated as

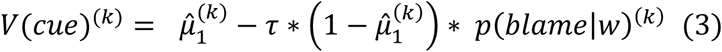

where 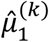 indicates p(correct|cue), and p(blame|w) has a negative value. Note that to transform p(blame|w) into a negative value, the probability of an incorrect cue 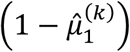 was multiplied by p(blame|w) because blame appears only when the choice is wrong, and the parameter *τ* represents the degree to which p(blame|w) negatively affected the total decision value computation and was bounded > 0 (because we assumed that the blame does not have a positive value). This parameter was also estimated in the log space with the prior means and variances of 0 and 1, respectively. Similarly, the total decision value of the other cue was calculated as follows:

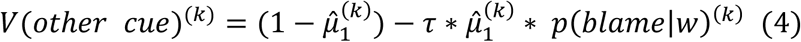

Note that 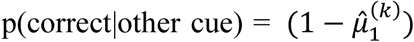 and 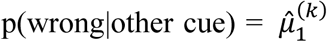 because of their reciprocity. Then, the total decision values of the two cues (*V*(*cue*)^(*k*)^ and *V*(*other cue*)^(*k*)^) were substituted for the original decision values of the cues within the RWS, HGF3, UDN, and CUDN response models (i.e., 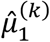 and 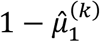) to make a decision, thereby generating four additional models: RWS-N, HGF3-N, UDN-N and CUDN-N. Importantly, the substitution of 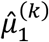 and 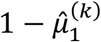 with *V*(*cue*)^(*k*)^ and *V*(*other cue*)^(*k*)^ in the Softmax model (equation [1]) modified the equation as follows:

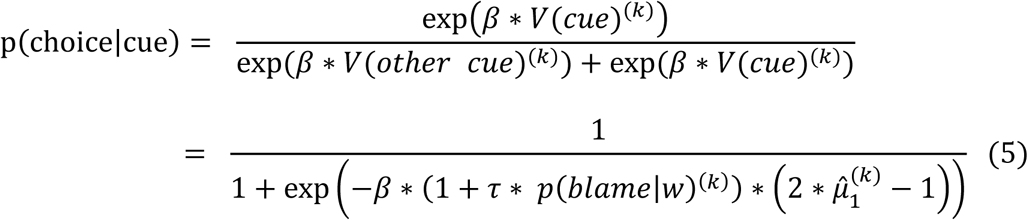

The term *β* * (1 + *τ* * *p*(*blame*|*w*)^(*k*)^) functions as the inverse temperature of the original Softmax model; thus, decision noise decreases as p(blame|w) increases. Consequently, in the RWS-N, HGF3-N and UDN-N models, p(blame|w) decreases decision noise, regardless of the uncertainty level (uncertainty-independent control of decision noise), which differs from the CUDN model in which p(blame|w) controls the uncertainty-induced decision noise but not the decision noise itself. Furthermore, in the CUDN-N model, p(blame|w) contributed both as a negative value and as the controller of the UDN, such that the β in equation (5) is the function of uncertainty and p(blame|w) as in the CUDN model.

In summary, eight models (RWS, HGF3S, UDN, RWS-N, HGF3S-N, UDN-N, CUDN-N and CUDN) were designed. The first three models did not consider the effect of p(blame|w) on the decision; the RWS-N, HGF3S-N, UDN-N models considered p(blame|w) as having a negative value and subsequently influencing decision noise regardless of uncertainty. Finally, in the CUDN model, p(blame|w) controls UDN, and in the CUDN-N model, p(blame|w) controls UDN and contributes as a negative value.

### Supporting Text 5. Priors of the parameters used in behavioral models

The prior means and variances of all parameters that were not newly introduced into our models were set to their default values in the TAPAS toolbox. Because we suspected that *v* might be unreliably estimated in the CUDN model given the large correlation between *v* and other parameters due to the interaction with *ζ* (because these factors are multiplicative), we used a narrow prior variance of 0.5 with a mean of 0 to ensure that the value of this parameter was close to 0 (if it was set to 0, then p[blame|w] had no effect on behavior, S2 Table). Similarly, *ζ* was bounded > 0; if the signs of both parameters (*v* and *ζ*) were not fixed as positive or negative, then we would not be able to determine them (e.g., if *ζ*\**v* = 1, then both [*ζ* = 1, *v* = 1] and [*ζ* = −1, *v* = −1] would be possible values for the [*ζ, v*]). Based on a previous study showing that uncertainty increases decision noise (Gershman, 2018), we bounded *ζ* > 0 and varied the sign of *v*.

### Supporting Text 6. Parameter correlation

We tested whether the parameters of the CUDN model were correlated. Two significant correlations were identified between these parameters, including a correlation between *ζ* and *β*_0_ (S1 Fig, Spearman’s ρ = 0.42, p = 0.004) as well as between *ζ* and ω (S1 Fig, Spearman’s ρ = −0.35, p = 0.018). Notably, no parameters were correlated with the parameter of interest, *v* (S1 Fig).

### Supporting Text 7. Parameter and behavioral pattern recovery simulation

We conducted a parameter recovery analysis by simulating the responses 100 times using the estimated parameters of every participant and re-estimating parameters for these responses to determine whether the parameters of the CUDN model were reliably estimated. Then, we performed a correlation analysis between the original parameters and the mean of the recovered parameters simulated 100 times. The results showed that all of the original parameters were highly correlated with the mean of the recovered parameters (all r > 0.75, all p < 10^−9^). Therefore, we concluded that the CUDN model reliably estimated all of the parameters. Furthermore, the simulated responses using the estimated CUDN model parameters of each replicated participant response significantly increased choice accuracy in high-blame blocks compared with the low-blame blocks (mean accuracy = 0.582 vs 0.566, t[4599] = 21.23, p < 0.001, CIs: 0.01 to 0.02, *d* = 0.3; two-tailed paired *t*-test, Fig 2D, left panel). Moreover, another simulation conducted with the increased value of the parameter *v* (+ 0.3) from each participant’s model showed a significantly greater increase in accuracy compared with the simulation using the original *v* (increase in accuracy: 1.6% before increasing *v* vs 1.9% after increasing *v*, t[4599] = 3.85, p < 0.001, CIs: 0.2% to 0.6%, *d* = 0.06; two-tailed paired *t*-test, Fig 2D, right panel), indicating that an increase in the degree of UDN suppression enabled participants to make more accurate choices during the high-blame blocks.

### Supporting Text 8. Comparison between depressed participants and the non-depressed participants

Of the 46 participants, eight were diagnosed with major depressive disorder (MDD). We compared the two groups with regard to task accuracy and the number of missing trials using two-sample t-tests to determine whether significant group differences existed in the performance on the experimental task between participants with and without depression. No differences in accuracy (mean accuracy = 0.57 [non-depressed] vs 0.587 [depressed], t(44) = − 0.97, p = 0.339, CIs: −0.05 to 0.02, d = −0.4) or number of missing trials (mean number of missing trials = 2.9 [non-depressed] vs 2.5 [depressed], t(44) = 0.24, p = 0.339, CIs: −3.1 to 4, d = 0.1) were observed. Furthermore, we also compared the differences in the CUDN model parameters between participants with and without depression using two-sample t-tests and found no differences between the two groups (all p > 0.2).

### Supporting Text 9. MRI acquisition

MRI acquisition was performed using a Siemens Verio Syngo 3T MR scanner (Siemens Healthcare, Erlangen, Germany). A total of 844 functional MRI image volumes were acquired using a T2 * 2D multiband echo-planar imaging (EPI) sequence (repetition time [TR] = 1500 ms, echo time [TE] = 32, flip angle = 50°, voxel size = 2 × 2 × 2 mm^3^, FOV = 225 mm, and number of slices = 64). A field map was also acquired for distortion correction. During the fMRI data acquisition, respiration and pulse data were acquired using a breathing belt and pulse oximetry for all participants but one due to a technical problem (YAD_10032). A high-resolution structural image was obtained using a T1-weighed sequence (TI = 1000 ms, TR = 2400 ms, TE = 2.02 ms, flip angle = 8°, voxel size = 1 × 1 × 1 mm^3^, and FOV = 224 mm).

### Supporting Text 10. fMRI preprocessing

All fMRI data preprocessing and analyses were performed using Statistical Parametric Mapping 12 (SPM12) (Penny, Friston, Ashburner, Kiebel, & Nichols, 2011). During preprocessing, spatial realignment, unwarping using individual field maps, normalization to the Montreal Neurology Institute (MNI) space and smoothing with a 5-mm full-width at half maximum (FWHM) Gaussian kernel were applied to the raw functional images. Additionally, 18 physiological noise regressors were computed using the PhysIO toolbox (Kasper et al., 2017), which is part of the TAPAS toolbox. Therefore, 24 regressors of no interest, including 6 motion-related regressors from the SPM preprocessing step (realignment) and 18 physiological noise regressors, were generated. Notably, we failed to obtain physiological data in one participant (YAD_10032) because of a physiological recording hardware problem. Therefore, we included only 6 motion-related regressors of no interest from the SPM preprocessing step for this participant.

### Supporting Text 11. Additional results and discussion of the parametric modulation analysis

The regions activated by BPE included the anterior and middle cingulate cortices cluster (ACC/MCC, [−6, −20, 32] in the MCC [−4, 26, 26] and ACC, cluster-level FWE-corrected p < 0.001, S5 Table and Fig 3) and the periaqueductal gray (PAG, [4, −30, −6], cluster-level FWE-corrected p < 0.001, S5 Table and Fig 3), which are related to a reactive fear circuit (Mobbs et al., 2007; Qi et al., 2018). In particular, the PAG signals an aversive prediction error (APE) induced by physical pain (Roy et al., 2014). Except for the rlPFC, large clusters, including the posterior portion of the mPFC including dACC, were activated by the suboptimal value (dmPFC, [6, 32, 40], cluster-level FWE-corrected p < 0.001, S6 Table and Fig 2). Activity in this region is associated with the encoding of conflict (Cavanagh et al., 2011) and differed from the activation pattern of p(blame|w), activated an anterior portion of the mPFC. These regions activated by a suboptimal value, including the rlPFC, dACC and IPL, might be related to the encoding of a suboptimal value, although it remains possible that these regions are related to other processes that increase in proportion to the suboptimal value, such as an outcome uncertainty (i.e., a variance in the Bernoulli distribution based on p[correct|cue]). However, considering the structure of the CUDN model and the result of the DCM/PPI analysis, the right rlPFC is likely to be involved in the calculation of a suboptimal value or the suboptimal value-related decision variables that lead to a suboptimal choice rather than the encoding of an outcome uncertainty that does not directly influence a suboptimal choice in our model. The roles that regions other than the right rlPFC play remain unclear.

### Supporting Text 12. PPI analysis using the PAG as an ROI and BPE as a psychological factor

Regions activated by BPE included the PAG. PAG signals an APE signals induced by physical pain, and these signals are conveyed to the anterior middle cingulate cortex (aMCC) (Roy et al., 2014). To determine whether BPE signaling is similar to the APE signaling induced by physical pain, we performed a PPI analysis using the PAG ROI. The PAG ROI was defined using the Harvard Ascending Arousal Network Atlas (Edlow et al., 2012). The mean BOLD signal time series of the PAG ROI was used as a physiological factor, and BPE was used as a parametric psychological factor. BPE positively modulated the functional connectivity between the PAG and aMCC ([8, 18, 40], cluster-level FWE-corrected p = 0.046 using a CDT of p < 0.05, S9 Table and S4 Fig). In addition to the neural and behavioral mechanism underlying UDN suppression, the prediction error signal produced by unexpected blame feedback involved the PAG and bilateral ACC/MCC. In particular, the PAG signals the APE induced by physical pain (Roy et al., 2014). In a previous study, the APE signal was conveyed to the aMCC (Roy et al., 2014). Similar to the APE signal in that previous study, the PPI analysis using the PAG ROI showed that the connectivity between the PAG and aMCC area increased with BPE (S9 Table and S4 Fig). These results suggest that the updating mechanism of blame expectation due to BPE is similar to that of physical pain expectation due to APE.

### Supporting Text 13. Dissociation of the neural activity between surprise and BPE

We also suspected that the result of the parametric modulation of BPE reflected the amount of surprise at an outcome, which is unrelated to blame. To test this possibility, we conducted an additional GLM that added surprise at the outcome to the original GLM, where surprise at the current trial was defined as the negative logarithm of the outcome of the current trial (Lawson, Mathys, & Rees, 2017), which increased as the likelihood of the current outcome (correct or incorrect) decreased. Importantly, in this additional GLM, we orthogonalized the regressors, and BPE was selected as the last parametric modulator at the outcome to remove the proportion of variance in BPE that might be explained by surprise at the outcome. Even after removing the shared variance with surprise, the activated area due to BPE was nearly identical to that of the original GLM. We then compared the activation pattern between BPE and surprise. Overlapping regions were observed in the dACC/aMCC and bilateral insula. Although the BPE activated the bilateral amygdala and adjacent temporal lobes, surprise did not activate the amygdala. Finally, surprise generated a large striatal activation, which is consistent with many previous reinforcement learning studies, while BPE did not. This result suggests that both BPE and surprise signal a saliency that involves the salience network (Uddin, 2015), while BPE is more likely to convey emotional information involving the amygdala, and surprise is more likely to convey a reward-related reinforcement signal that typically involves the striatum.

